# Correction of multiple-blinking artefacts in photoactivated localisation microscopy

**DOI:** 10.1101/2021.03.24.436128

**Authors:** Louis G Jensen, Tjun Yee Hoh, David J Williamson, Juliette Griffié, Daniel Sage, Patrick Rubin-Delanchy, Dylan M Owen

## Abstract

Photoactivated localisation microscopy (PALM) produces an array of localisation coordinates by means of photoactivatable fluorescent proteins. However, observations are subject to fluorophore multiple-blinking and each protein is included in the dataset an unknown number of times at different positions, due to localisation error. This causes artificial clustering to be observed in the data. We present a workflow using calibration-free estimation of blinking dynamics and model-based clustering, to produce a corrected set of localisation coordinates now representing the true underlying fluorophore locations with enhanced localisation precision. These can be reliably tested for spatial randomness or analysed by other clustering approaches, and previously inestimable descriptors such as the absolute number of fluorophores per cluster are now quantifiable, which we validate with simulated data. Using experimental data, we confirm that the adaptor protein, LAT, is clustered at the T cell immunological synapse, with its nanoscale clustering properties depending on location and intracellular phosphorylatable tyrosine residues.

## Introduction

Single molecule localisation microscopy (SMLM) methods, such as PALM, circumvent the diffraction limit of light by separating fluorophore detections in time through stochastic activation and photobleaching, and then localizing the resulting sparse distribution of point spread functions^1^. The resulting point-pattern is a purported realisation of the underlying ground truth positions of the fluorophores, but is corrupted by a number of artefacts resulting from the photophysical behaviour of the probes as well as the imaging and localisation steps. Most problematic is the multiple appearance (multiple-blinking) problem where fluorophores undergo multiple on-off cycles before permanently bleaching, combined with the discretization effects that result from observing fluorescent signals on discrete camera frames^2^. The multiple-blinking problem results in data sets that are artificially clustered and overly populated (Figure 1a). As such, quantitative cluster analysis of SMLM data, in particular testing for spatial randomness of the underlying fluorophores, remains a challenge.

**Figure 1:**
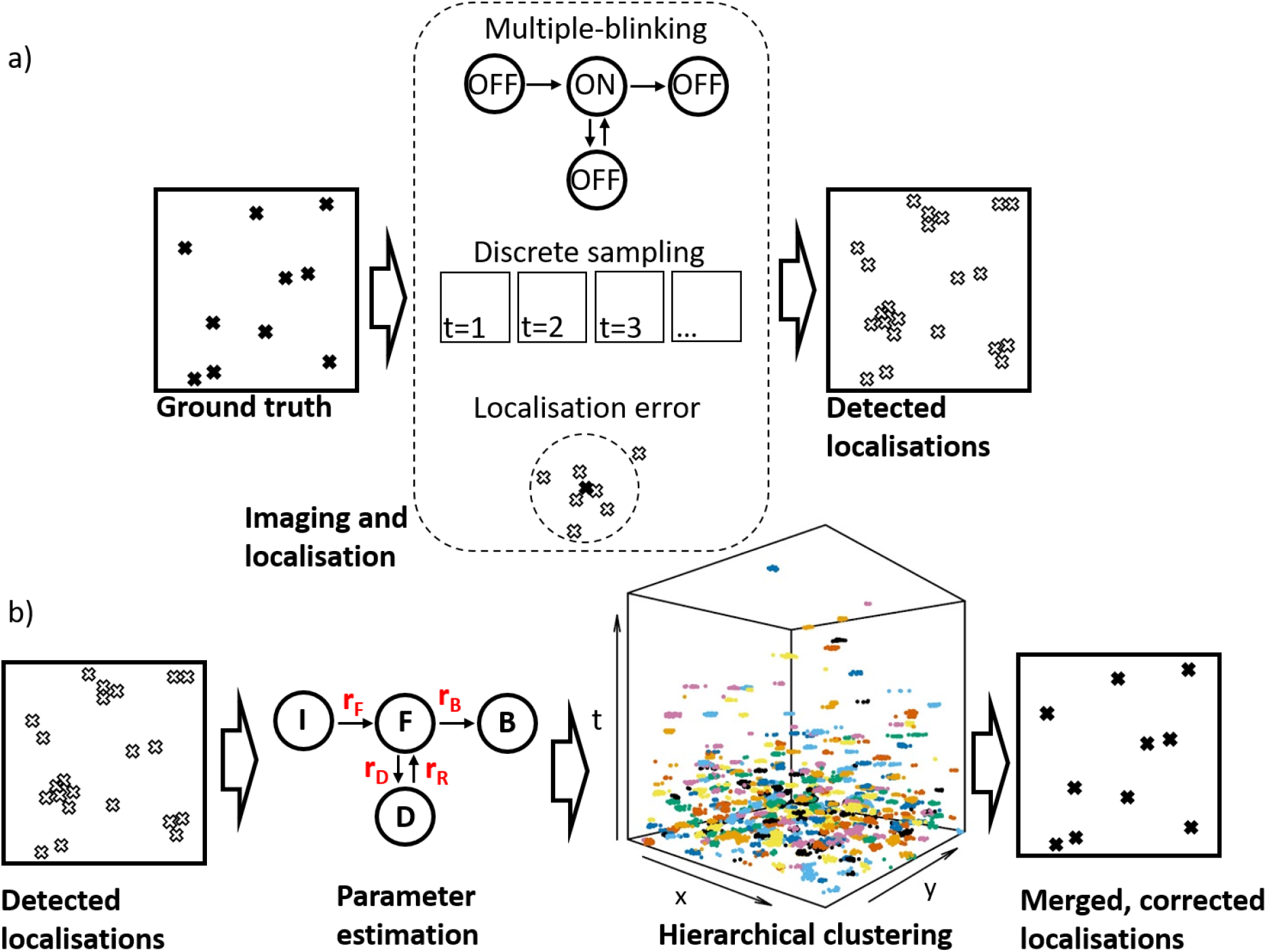
Illustration of the MBC workflow. a) During PALM image acquisition and subsequent localisation steps, the ground-truth protein positions are corrupted by multiple-blinking in combination with discretisation by the camera frames and scrambling by the localisation uncertainty, resulting in a data set which is over-populated and over-clustered. b) Our algorithm (MBC) takes as input x,y,t,σ data and estimates the rate parameters of a 4-state photophysical model, from which it derives the total number of molecules in the ROI. This is then used as input to a hierarchical clustering step (experimental data shown with colours representing the clusters found), after which clusters are merged to their centres, creating a new dataset free from multiple-blinking and with enhanced localisation precision.

The most commonly employed method for correction of the multiple-blinking problem is to merge events that appear close in space and time^3,4,5^. Such methods require a means of determining the best spatial and temporal thresholds for merging. This determination typically relies on heuristic methods, since the blinking behaviour of the fluorescent probes is often unknown, and can vary between experiments. Apart from the challenges involved in determining optimal thresholds, these methods have variable performance, depending on the underlying protein organization and blinking characteristics. Instead of attempting to produce a corrected version of the data which can then be used for any subsequent analysis, other approaches have looked to correct specific spatial statistics to account for multiple-blinking. For example, it is possible to use calibration data to estimate a multiple-blink corrected pair-correlation curve^6,7^. However this cannot then be used to find a cluster map, or count absolute numbers of molecules.

In this work, we present a new method for correction of multiple-blinking artefacts in PALM data, which estimates, directly from the sample data set, the parameters of a realistic model of fluorescent protein photophysics^8^. Cluster analysis of the spatio-temporal (x,y,t,σ) data set then allows computation of the marginal likelihood of any given blink-merge proposal, under a full generative model for the data. We select the most likely of several proposals generated using a customised hierarchical clustering algorithm. Finally, each blink cluster is consolidated into a single position, now free from multiple-blinking and with improved localisation precision. The overall effect is to convert the set of raw x,y,t,σ localisation data into a new set, x,y,σ, with enhanced resolution.

We validate the method on simulated PALM data, varying both the ground-truth organization (regular, random, clustered) and photophysical properties of the fluorescent proteins (light and heavy multiple-blinking). In each case, we compare to the state-of-the-art method of dark time thresholding (DTT). Our method allows for testing the completely spatially random (CSR) hypothesis at the correct significance level, whereas the thresholding method fails to do so, and also outperforms the state-of-art in every other metric (including ground truth recovery and extracted cluster properties).

PALM is increasingly used in the biological sciences and owing to the properties of commonly used total internal reflection fluorescence (TIRF) illumination, the distributions of membrane proteins have been especially well studied. Despite this, because of artificial clustering resulting from multiple-blinking, the question of whether membrane proteins are randomly distributed or not has become increasingly contentious^9^. Using our validated method combined with subsequent testing of the corrected protein locations, we show that the adaptor protein Linker for Activation of T cells (LAT) is clustered in the plasma membrane of CD4+ Helper T cell lines after the formation of an artificial immunological synapse^10,11^ against an activating, antibody-coated surface. However, subsequent Bayesian cluster analysis^12,13^ shows the clustering properties to be dependent on its macro-scale location within the synapse and on the presence of intracellular phosphorylatable tyrosine residues which mediate protein binding. We now propose that PALM, combined with the method we present here, can be used to test for spatial randomness in other membrane protein species.

## Results

### Description of the algorithm

We work with the space-time localisations and uncertainties that result from localisation software (here ThunderSTORM^14^) that is run on the raw microscope data. We apply drift correction, but otherwise no pre-processing is used. The data points are then modeled as a collection of independent and identically distributed fluorophore blinking clusters, with times following a realistic 4-state model^15,16^, discretized by the camera frames. The spatial locations for each cluster are independently drawn from a circular Gaussian distribution of fixed centre (the true molecule position) and variable but known standard deviation (the localisation uncertainty). The centres are given a uniform prior over the region of interest (ROI).

We refer to our algorithm as model-based correction (MBC), and a schematic of its workflow is shown in Figure 1b. We first estimate the temporal rates governing the switching behaviour of fluorescent proteins under the 4-state model^8^, and the fraction of background noise points. This is done directly using the experimental data, requiring no additional calibration experiments. A recently developed mathematical technique, (updated to remove one tunable parameter - see Online Methods), extracts a component from the empirical mark and pair correlation functions which depends only on the spatio-temporal dynamics of the multiple-blinking process, and not the underlying protein distribution. The parameters of the 4-state model drive the theoretical shape of this component, and so they can be optimised to best fit the empirical version^8^. The rate-estimates allow computation of the likelihood of a sequence of timepoints purported to correspond to one multiple-blinking fluorescent protein, and further yields an estimate on the total number, *N*, of proteins and noise points in the ROI. Using a custom agglomerative hierarchical clustering (HC) algorithm^17^, we split the data in the ROI into partitions with *N* categories. HC takes as input a dissimilarity matrix and a linkage criterion. The dissimilarity matrix determines the distances between pairs of points, and the linkage criterion determines the way to generalise this distance to pairs of clusters. To favour groups likely to correspond to multiple-blinking clusters, we first scale the temporal dimension by a time-dilation hyperparameter, *S*, and then compute the sum of Euclidean distances in space and in time. For linkage, we choose Ward’s Minimum Variance Method^19^, which is well-suited for Gaussian clusters, and consistently resulted in the most likely partitions across all tested linkage criteria. By varying *S*, we obtain a large sequence of blinking cluster proposals, and evaluate the marginal likelihood of each using a uniform prior on the partitions. Finally, using the best found partition and the localisation uncertainties, we optimally merge the clusters down to their estimated centres, using inverse-variance weighted averages, and update the uncertainty associated with that centre.

### PALM data simulation setup

For a given set of protein positions, corresponding PALM data were generated as follows. We simulated fluorescent protein time traces according to the 4-state switching model (see Figure 1b), and the continuous signals were discretised to emulate a camera operating at 25 frames per second (40 ms integration time). This was done for 2 different sets of rates (given in Table 1, Online Methods), with the light blinking resulting in 5.36 appearances per protein, and the heavy blinking resulting in 14.94 appearances. For each of these appearances, the observed spatial coordinates were simulated by adding Gaussian localisation noise to the ground-truth position of the associated fluorescent protein, with standard deviation following a Gamma distribution with mean 30 nm and standard deviation 13.4 nm, emulating the localisation uncertainties that can be observed in real PALM data^10^. Each simulated ROI was corrected using MBC and compared to correction using DTT (Figure 2a). For DTT, points were considered to have come from the same fluorophore if they were separated by at most 4 times the mean localisation uncertainty in space, and were no further than *T* apart in time, where the optimal *T* was determined for each ROI using the method of Annibale et al^4^.

**Figure 2:**
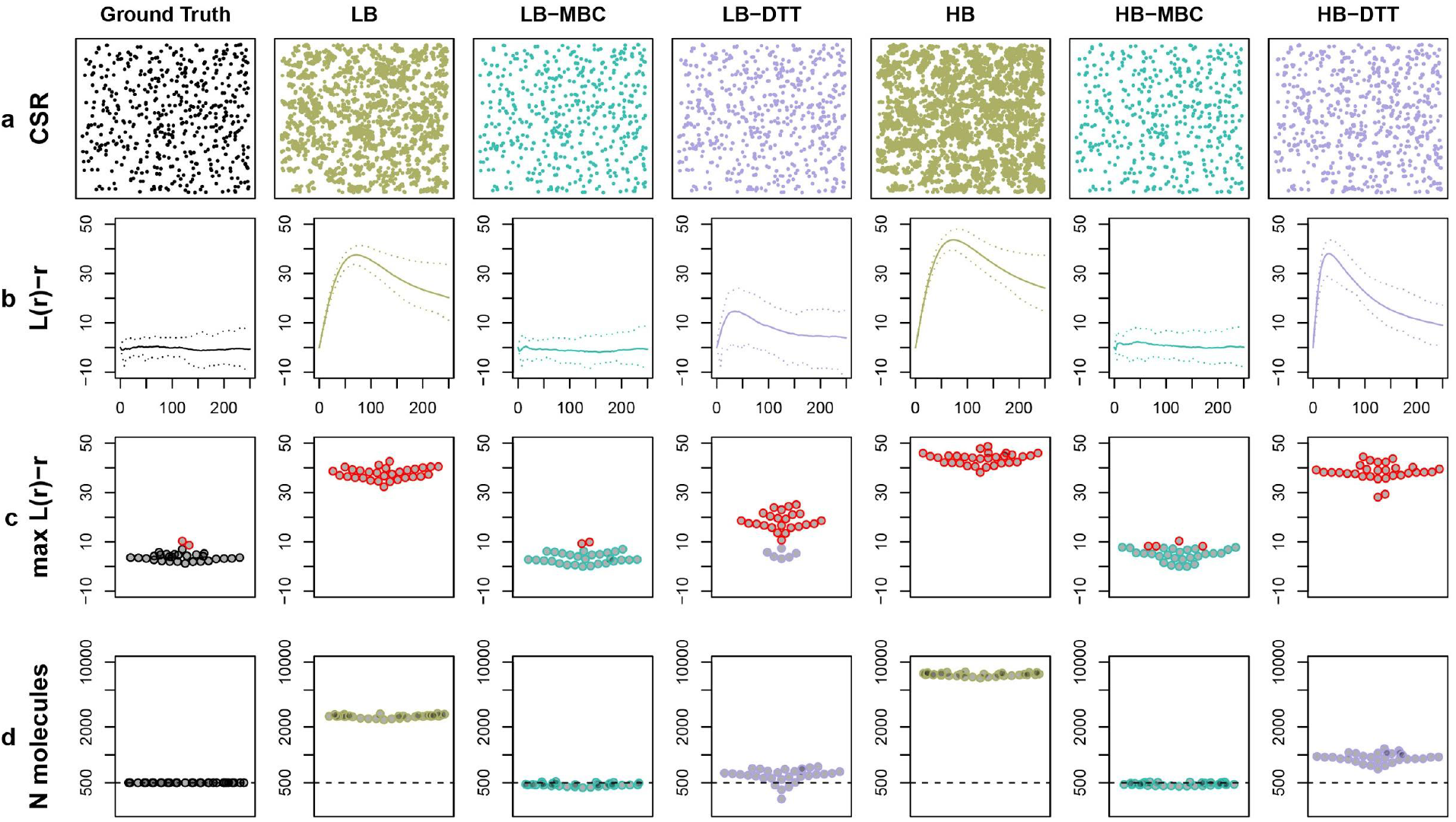
Testing for spatial randomness. a) Representative simulated data of ground-truth CSR points with light or heavy blinking (LB or HB) either corrected by MBC or DTT as a comparison. b) L(r)-r (mean in solid line) with pointwise 95% quantile bands (dashed line). c) max(L(r)-r) derived from these functions. Points in red correspond to ROIs that were rejected as CSR in a Monte-Carlo test (p <= 0.05). Note that DTT often (and sometimes always) incorrectly rejects the CSR null hypothesis, whereas MBC does not. d) Number of molecules per ROI (log-scaled) showing superior correction of MBC compared to DDT in light and heavy blinking cases.

### Testing for complete spatial randomness

We first evaluate our algorithm for testing for complete spatial randomness of the underlying ground-truth proteins. In each run (*n =* 30 per condition), 500 proteins were placed at random in a noiseless 3000 nm x 3000 nm ROI. For each ROI, we compute the function *L(r)-r* (Figure 2b), where *L* is Besag’s *L* function^19^, testing its maximum (Figure 2c) under a CSR null hypothesis. The standard DTT correction method was unable to recover the ground-truth functions and resulted in rejection of the CSR null hypothesis in 24 and 30 out of the 30 regions, for light and heavy blinking respectively. On the other hand, MBC resulted in the CSR null hypothesis being rejected for 2 and 4 of the regions for light and heavy blinking respectively. These numbers are within the expected range at a 5% confidence level. Thus, we were able to reliably test the CSR hypothesis using MBC, but not using DTT. The estimated total number of fluorescent proteins in each ROI is shown in Figure 2d. Under CSR, DTT tends to overestimate the number of proteins in the ROI whereas MBC closely recovers the ground truth.

### Cluster analysis

In this experiment, we demonstrate that a clustering algorithm can extract correct cluster descriptions from underlying clustered ground truth protein distributions when coupled with MBC, and we compare performance with DTT. We simulated data from 2 clustered protein distributions (n = 30 per condition). In each run, 500 ground-truth proteins were placed in a 3000 nm x 3000 nm ROI, with either 10 clusters of 10 molecules each, overlaid with 400 CSR molecules (light clustering) or 10 clusters of 40 molecules each, with 100 overlaid CSR molecules (heavy clustering). Clustered points were simulated as symmetric Gaussian clusters with a standard deviation of 30 nm, and the cluster centres were uniformly distributed over the ROI. Again, both light and heavy blinking was then added to the localisations, resulting in 4 conditions of spatial organisation and blinking characteristics (Figure 3a). We used Bayesian cluster analysis^12^ for detection of clusters in MBC and DTT corrected data sets. Only MBC could consistently recover the 10 clusters under varying degrees of blinking severity (Figure 3b). The failure of DTT to recover the correct number of clusters is even more evident in the case of heavy clustering (Figures 3c and d).

**Figure 3:**
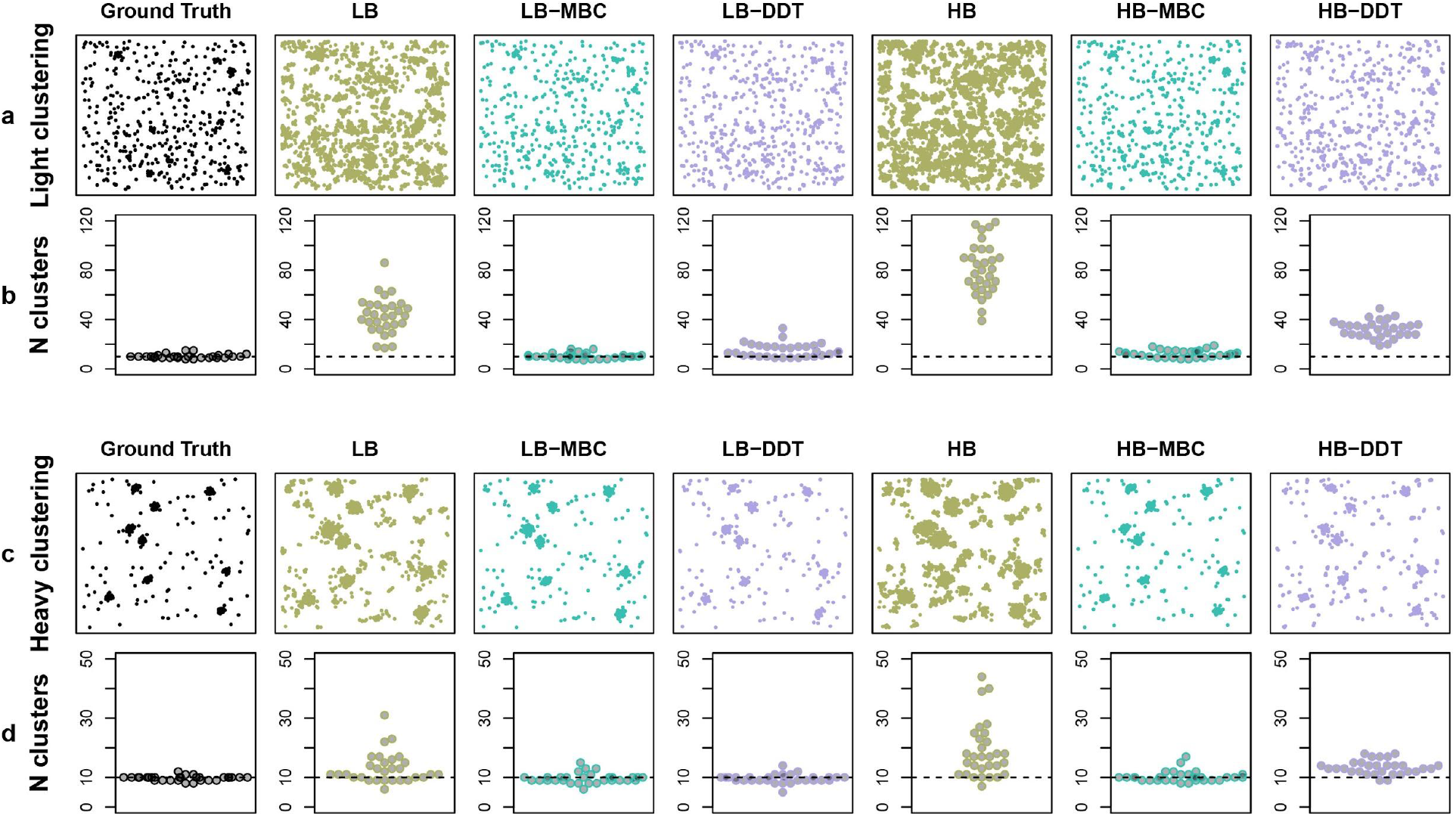
Testing on clustered ground-truth data sets. a) Low levels of clustering with either light or heavy blinking, corrected by MBC or DDT. b) Number of detected clusters (true number of clusters in dashed line) by Bayesian analysis. c) High levels of clustering with either light or heavy blinking, corrected by MBC or DDT. d) Number of detected clusters (true number of clusters in dashed line) by Bayesian analysis. MBC has superior performance in all cases except heavy clustering/light blinking, where results are comparable.

### Recovery of the ground truth

In addition to simulating realistic data, we also consider a more controlled, synthetic setup wherein ~500 fluorescent proteins are regularly positioned on a ~3000 nm x 3000 nm grid. The nature of this dataset allows for easier comparative visualisation of the performance of MBC and DTT (Figure 4a), and the improvements offered by our method are visually clear. To validate this, we also compute the 1st Wasserstein distance between the true and corrected grids. This can be thought of as the cost of transporting a standardised mass between two sets of points, and is also known as the earth mover’s distance. For a perfectly reconstructed grid, this distance is zero, with any discrepancy increasing the distance. For 50 realisations of both light and heavy blinking on the grid, we see that MBC presents an improvement over DTT, with a lower distance to the true grid, particularly under heavy blinking conditions (Figure 4b). Because MBC is robust to heavy blinking, relative to DTT, it is in fact able to use heavy blinking to its advantage, by averaging localisation precisions from large groups of merged observations (Figure 4c).

**Figure 4:**
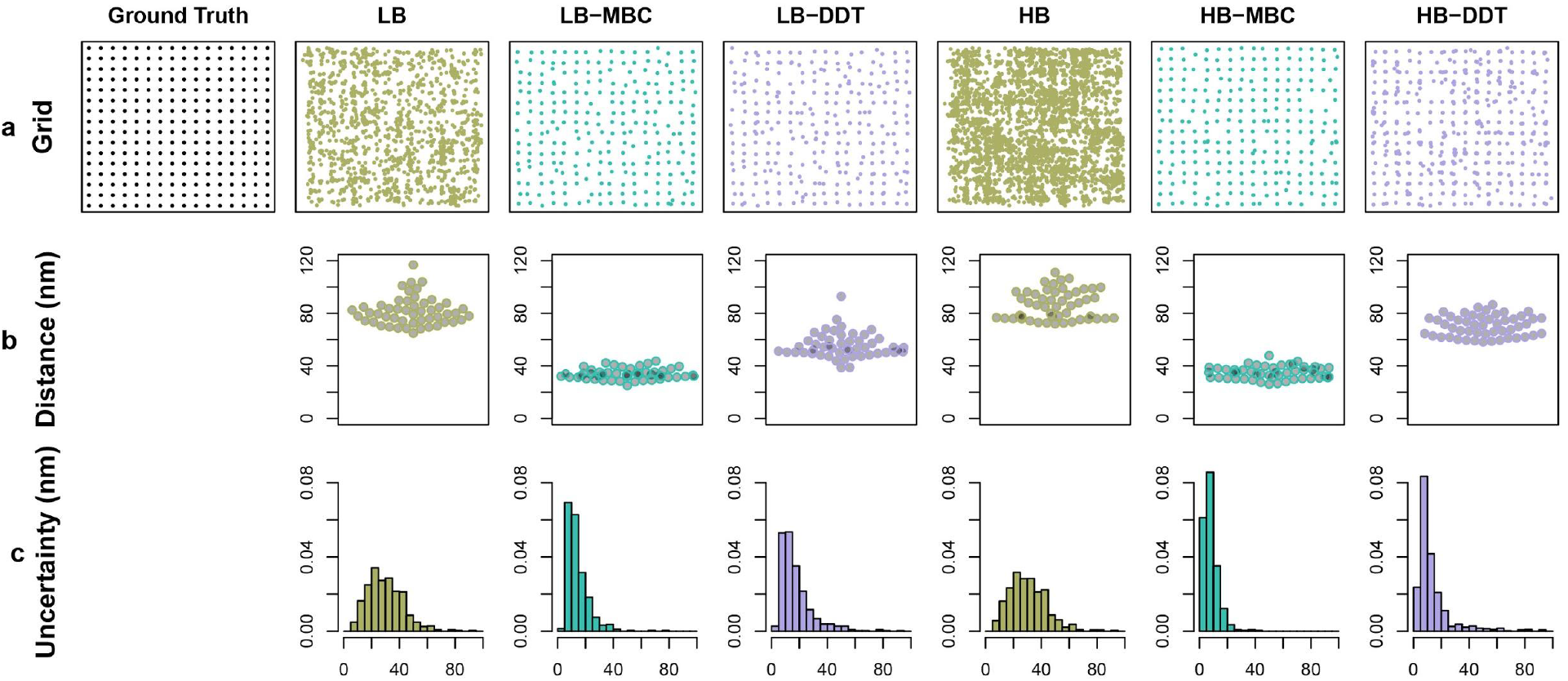
Testing against molecules on a fixed grid. a) Ground truth and representative simulated data. b) Wasserstein distances between simulated data and ground truth showing that MBC generates output closer to the ground truth. c) Normalised histograms of localisation uncertainties of individual molecules (nm) showing that MBC also generates increased localisation precisions compared to uncorrected data or DDT.

### Determining optimal imaging conditions

As a final test using known ground-truth simulated data, we used Virtual-SMLM^20^ to simulate raw camera frames. This allowed us to test the effect of varying both the camera frame rate and the intensity of the 405 nm activation laser on the performance of MBC. A ground truth of CSR fluorescent proteins were simulated (Supplementary Figure 2a), imaged using the virtual microscope and output analysed with ThunderSTORM. The camera integration time was set to either 10 ms or 40 ms and the 405 nm laser intensity either kept constant, or ramped to maintain a constant density of point spread functions (PSFs) per frame over the course of the acquisition. Raw localisations (Supplementary Figure 2a) were then corrected using MBC (Supplementary Figure 2b). The Wasserstein distance shows marginally superior performance of the reconstruction when using constant 405 laser power and when using longer, 40 ms frames. We attribute this to the lower density of PSFs per frame in the constant-405 case leading to fewer overlapped PSFs during localisation and to the increased localisation precision offered by the longer frames (Supplementary Figure 2c). The performance of MBC itself is only weakly dependent on the imaging conditions, and in each condition we were able to recover the ground truth number of molecules to within around 10% error. We conclude therefore that when using MBC, PALM imaging conditions should be chosen to maximise conventional notions of data quality — low density of PSFs and high signal-to-noise ratio. Because of this, we also conclude that MBC is also backwards compatible with all historically acquired PALM data.

### Analysis of experimental data

Nanoscale clustering is posited to play a role in regulating protein-protein interactions and therefore the efficiency of signalling propagation along pathways^21^. In the context of an immune response, T cell microclusters of proximal signalling molecules have been widely documented by conventional total internal reflection fluorescence (TIRF) microscopy^22,23^. Many of these have recently been studied by SMLM and shown to also be clustered on the nanoscale^10,11,24,25^. The claim has proved controversial however, with counter-proposals that, in some circumstances, proteins may in fact be randomly distributed on the cell surface, with observed clustering attributed to multiple-blinking artefacts inherent to SMLM^9^. For PALM data, MBC should enable researchers to navigate this controversy.

To demonstrate the application of MBC to experimental data, we analysed the distribution of the adaptor protein LAT^26^ in the plasma membrane of the Jurkat CD4+ Helper T cell line at an artificial immune synapse formed against an activating, antibody coated coverslip (see Online Methods). To assess the role of intracellular phosphorylation in maintaining this distribution, we also mutated intracellular tyrosine residues to phenylalanine (YF LAT). Both wild-type (WT) LAT and YF LAT were fused to the photoconvertible fluorescent protein mEos3.2 with cells imaged under TIRF illumination. Raw localisations were obtained using ThunderSTORM and then corrected using MBC. The resulting corrected localisations were then tested for spatial randomness using the L-function, and any regions found to be clustered subjected to Bayesian cluster analysis^12^.

Figure 4 shows WT and YF LAT-mEos3.2 from representative regions acquired from the central regions of the cell synapse and from the synapse periphery, both before (Figure 4a) and after (Figure 4b) correction using MBC. Clearly, the large, dense clusters evident in the uncorrected data in all conditions are reduced in the corrected regions. However, by analysing the L-function curves from the ROIs (Figure 5c) and extracting the maximum value of those curves (Figure 5d), we were able to perform significance testing on whether the LAT distributions were truly CSR. For the two WT LAT conditions, the null hypothesis that LAT is randomly distributed was rejected in most regions. Therefore, it is likely that WT LAT was clustered in most analysed WT ROIs. This was not true for the YF mutant however, with the null hypothesis of randomly distributed LAT not rejected in the majority of peripheral regions (Figure 5d). This therefore may point to a role of intracellular tyrosine phosphorylation in maintaining LAT clustering.

**Figure 5:**
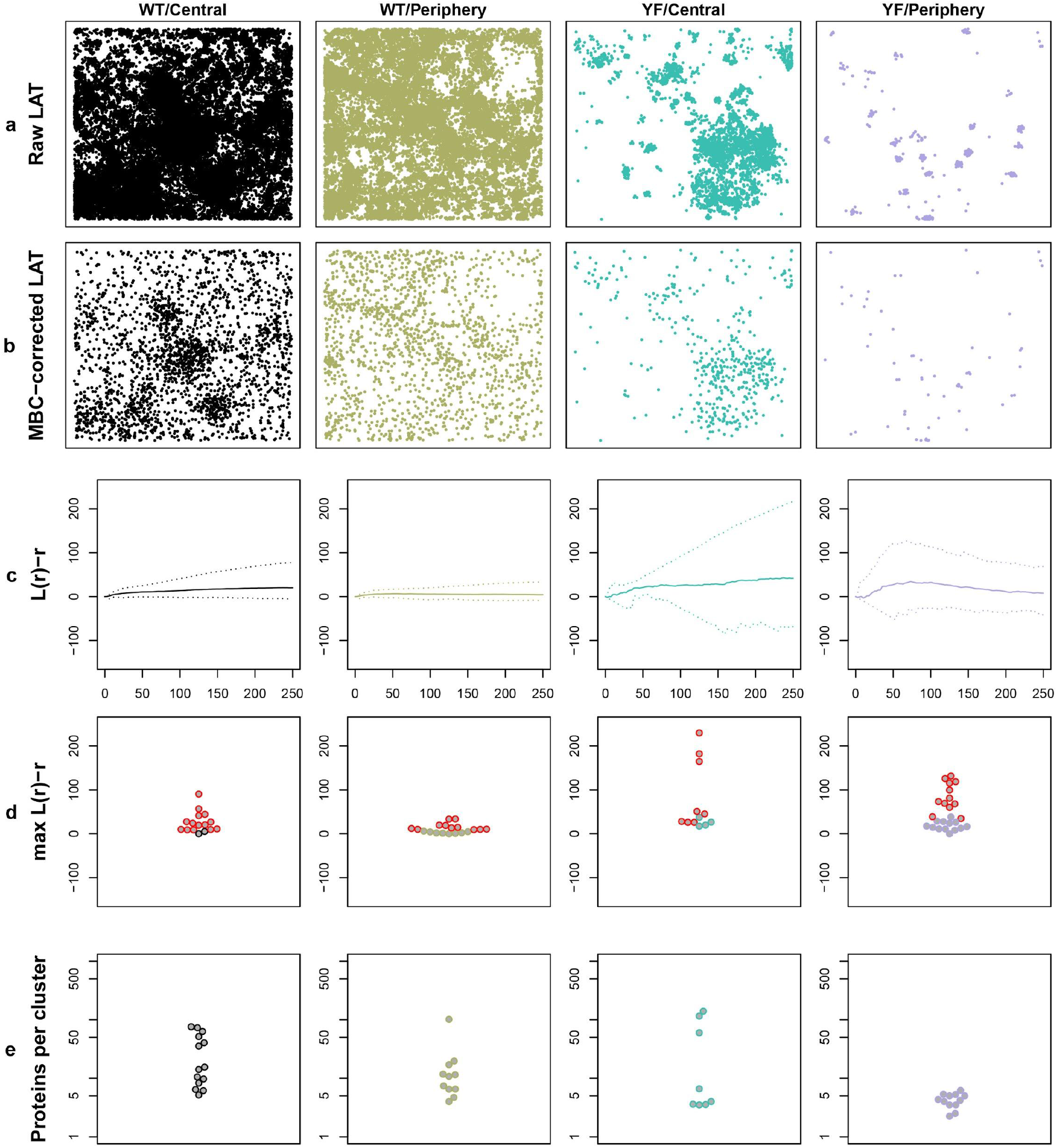
Cluster analysis of LAT-mEos3.2 at the T cell immunological synapse. a) Representative, raw 3000 nm x 3000 nm ROIs from each of the 4 conditions (WT Centre, WT Periphery, YF Centre and YF Periphery). b) Representative MBC-corrected ROIs, on which analysis was conducted. c) L(r)-r (mean in solid line) with pointwise 95% quantile bands (dashed line). d) max(L(r)-r) derived from these functions. Points coloured in red correspond to ROIs where the CSR null hypothesis was rejected in a Monte Carlo test (p < 0.05). These ROI were then retained for subsequent Bayesian cluster analysis. e) Number of proteins per cluster detected by Bayesian analysis.

For all regions where the CSR null hypothesis was rejected (treated as clustered regions) we then further interrogated the data using Bayesian cluster analysis. In addition to existing cluster descriptors output by the algorithm, the number of points per cluster (cluster membership) is now biologically relevant, since this now represents the number of real fluorophores, not detected localisations. For WT LAT the data showed no statistically significant difference in cluster membership between central and peripheral regions. However, the YF mutant showed a significant decrease in the number of molecules per cluster in peripheral regions, both when compared to YF central regions (p = 0.026) and compared to WT peripheral regions (p = 0.001) (Figure 5e). Other outputs from the cluster analysis: number of clusters per ROI, cluster radius, percentage of molecules in clusters, total molecules per ROI and the relative density of molecules inside and outside clusters are shown in Supplementary Figure 2, with p-values summarised in Supplementary Table 1. The decrease in cluster membership and, in some ROIs, the loss of clustering altogether, in peripheral regions of the T cell synapse resulting from the YF mutation, is a strong indication that intracellular tyrosine phosphorylation is involved in maintaining LAT signalling clusters. Signalling phosphorylation events are known to originate in the synapse periphery and it is therefore consistent that the effect of the mutation is most pronounced there, compared to the central region where signalling is terminated^27^.

## Discussion

Super-resolution fluorescence microscopy by SMLM, such as PALM, results in a pointillist data representing an attempted realisation of the underlying ground-truth fluorophore locations^1^. A common goal in the biological sciences is to test whether such underlying distributions are clustered or randomly distributed and, if clustered, to determine their clustering properties. Achieving this goal has proved difficult however, because the generated localisations are corrupted by artefacts, principally the repeated localisation of the same fluorophore due to multiple-blinking^2^. This has led to controversy about whether proteins are truly clustered in cells, hindering our understanding of the causes and function of nanoscale protein clustering.

Here, we develop an algorithm, MBC, for correcting multiple-blinking that requires no user input, no additional calibration data, and is not limited to a specific analysis goal. We show that it can be used to reliably test for spatial randomness or recover other clustering properties from the ground truth. A number of methods have been put forward to test for spatial randomness in SMLM data. These include, for example, methods based on varying the labelling density and observing the effects on specific cluster analysis outputs^28^ or by labelling the same species with two different fluorophores allowing a cross-comparison to be made^29^. These methods, however, require multiple sample preparation rounds and are therefore more complex and time-consuming. Correction can also be made by measuring blinking behaviour in a separate sample of well-isolated fluorophores^7^, but this again adds complexity and experimental effort and requires the assumption that probe photophysics is maintained between the sample and the calibration. It is also possible to measure or simulate multiple-blinking using realistic photophysical models and use these to derive new CSR confidence intervals for the L-function curves^30^. It should be noted however that while all of these competing methods can be used to account for multiple-blinking, none produce a new set of corrected positions and therefore none can be used to extract rich descriptors such as those output by a clustering algorithm. MBC therefore represents a new capability - of obtaining a set of corrected ground-truth locations of sufficient quality that any subsequent statistical analysis can be conducted with assurance.

The limitations of MBC are as follows. The method is only applicable to the four-state photophysical model typical of PALM acquisitions, and therefore cannot be used to correct dSTORM or other SMLM modalities. Performance of the correction will decrease as the clustering of the ground-truth increases, however, it tested favourably with realistic and heavily clustered scenarios. The method also adds computational time to any analysis pipeline. For a 3000 nm x 3000 nm ROI containing 500 ground-truth proteins, we estimate the MBC step to take 3-5 minutes per ROI on a standard desktop computer. Of course, as it results in fewer points per ROI, subsequent analysis will typically be accelerated. While here the correction is limited to 2D data, it can in principle be adapted to 3D x,y,z coordinates.

In conclusion, MBC allows for accurate recovery of ground-truth fluorophore positions, with enhanced precision, from PALM data sets subjected to multiple-blinking artefacts. For the first time, these corrected sets are of sufficient quality to allow accurate cluster analysis and the statistical testing for complete spatial randomness. We therefore believe that PALM combined with MBC will be an invaluable tool for addressing questions on the existence, determinants and functions of protein nanoscale clustering.

## Supporting information

Code and example data

## Acknowledgements

L.G.J was supported by the Centre for Stochastic Geometry and Advanced Bioimaging, funded by grant 8721 from the Villum Foundation. D.O. acknowledges funding from BBSRC grant BB/R007365/1. P.R.D. acknowledges funding from the BBSRC grant BB/R007837/1. We also acknowledge the use of the King’s College London Nikon Imaging Centre (NIC). The authors would like to thank Ute Hahn (Aarhus University) for motivating this project in initial discussions.

## Author contributions

L.G.J. developed the software. L.G.J., T.Y.H., D.M.O. and P.R-D conceived the experiments. L.G.J. and T.Y.H. ran analysis. L.G.J. and T.Y.H. provided simulated data. D.J.W., J.G. and D.S. provided additional simulated and experimental data. L.G.J., T.Y.H., D.M.O. and P.R-D wrote the manuscript. L.G.J. and P.R-D. conceived the method.

## Online Methods

### Blinking simulation parameters

**Table 1:** Rate parameters used to simulate multiple-blinking for the light and heavy blinking cases, using the 4-state model shown in Figure 1B.

| *Rate parameter* | *Light Blinking (LB)* | *Heavy Blinking (HB)* |
| --- | --- | --- |
| $r_F$ | $0.005\text{ s}^{-1}$ | $0.005\text{ s}^{-1}$ |
| $r_B$ | $8\text{ s}^{-1}$ | $2.5\text{ s}^{-1}$ |
| $r_D$ | $10\text{ s}^{-1}$ | $10\text{ s}^{-1}$ |
| $r_R$ | $0.75\text{ s}^{-1}$ | $0.75\text{ s}^{-1}$ |

### Sample preparation

For LAT images, Jurkat E6.1 cells (ECACC 88042803) expressing LAT-mEos3.2 (wild-type, WT LAT, or signalling deficient mutant, YF LAT) were introduced to anti-CD3 (at 2 μg/ml; eBioscience clone OKT3, 16-0037-81) and anti-CD28 (at 5 μg/ml; RnD Systems, clone CD28.2, 16-0289-85) coated glass-bottomed chamber slides (#1.5 glass, ibidi μSlides) at 50 × 10^3^ cells/cm^2^ in warm HBSS and incubated at 37°C for 5 minutes to allow for synapse formation. The chamber wells were gently washed with warm HBSS and then fixed in 3% paraformaldehyde in phosphate-buffered saline (PBS) for 20 minutes at 37°C. Fixed cells were washed five times in PBS and used immediately for PALM imaging.

### Imaging

PALM image sequences were acquired on a Nikon N-STORM system in a TIRF configuration using a 100 × 1.49 NA CFI Apochromat TIRF objective for a pixel size of 160 nm. Samples were continuously illuminated with 561 nm laser light at approximately 2 kW/cm^2^ and 405 nm laser light (to induce photo-conversion) at approximately 2 W/cm^2^. Images were recorded on an Andor IXON Ultra 897 EMCCD with an electron multiplier gain of 200 and pre-amplifier gain profile 3 to a centered 256 × 256 pixel region at 40 ms per frame for 5,000 to 15,000 frames.

### Virtual microscope simulations

Raw camera frames were generated using Virtual-SMLM^20^ operating in PALM mode (i.e., using a 4 state photophysical model). The frame rate was set to 25 or 100 frames per second. The activation laser (i.e. initial state transition) was either fixed or ramped up over the acquisition. In the first case, the number of fluorophores emitting per frame decreases over time. In the second case, it remains constant over the acquisition. Emission traces were generated independently for each fluorophore and imaging continued until all fluorophores had been imaged and bleached. All other state transition probabilities and photophysics properties were fixed to mimic mEos blinking characteristics. The PSFs were recorded on a virtual EMCCD camera, with an EM gain fixed at 300. Virtual-SMLM took as input ground truth maps of mEos2 positions. 5556 mEos proteins were placed randomly over a 10000 nm x 10000 nm 2D area. Generated camera frames were then analysed using ThunderSTORM and the data cropped into non-overlapping 3000 x 3000 nm regions.

### Localisation

Localisation of fluorophore coordinates were reconstructed using ThunderSTORM^14^ and corrected for sample drift using cross-correlation of images from 5 bins at a magnification of 5. No further post-processing was performed.

### Mathematical details

#### 1. Marginal likelihood of clusters

We represent the observed process by a series of localisations 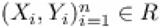 with associated ‘blink’ times *T*_1_,…,*T_n_*, and localisation uncertainties 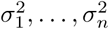, where *R* = [*x*_0_*,x*_1_] × [*y*_0_*,y*_1_] is the region of interest.

For a given partition of the localisations into groups, we compute the marginal likelihood of the data as follows. Consider a group comprising the observations 1*,…,m*, with 1 ≤ *m* ≤ *n* (without loss of generality), *posited to correspond to one, distinct, molecule*. In particular, we defer until later the treatment of background noise.

The independence assumptions set out in the main article result in the following marginal likelihood factorisation:

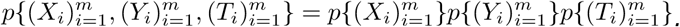

Denoting by *μ* = (*μ_X_,μ_Y_*) the true position of the molecule, the spatial components above have likelihood (given only for 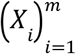)

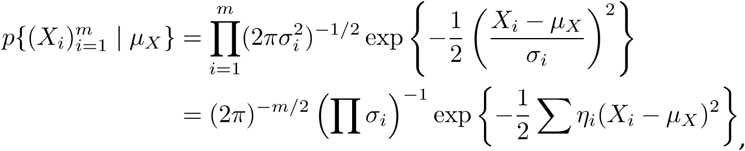

where 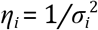. Defining the weighted mean

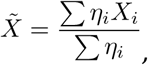

we find

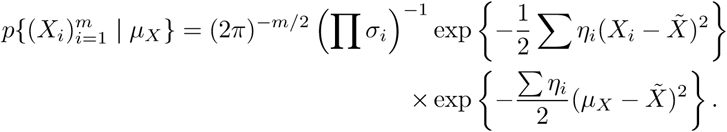

Placing a uniform prior on *μ_X_*, we find

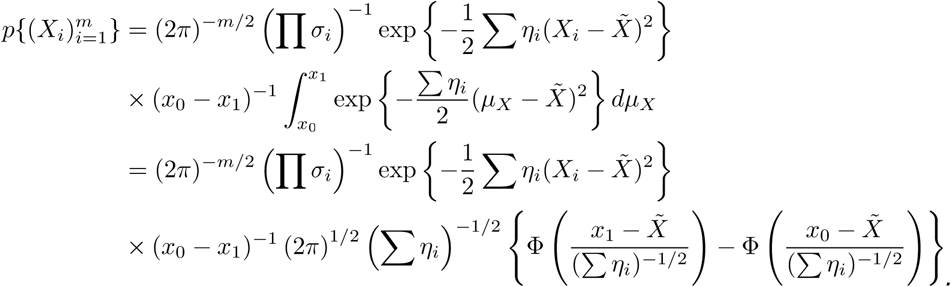

The temporal component has likelihood

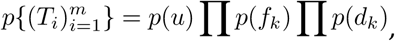

where each term is computed as follows. The blink times *T*_1_,…,*T_m_* are not typically observed exactly, and instead one has access only to associated frame numbers *F*_1_,…,*F_m_*, taken to represent (small) windows of time containing them. We therefore consider a visit to the fluorescent state to be a block of *L* ≥ 1 contiguous fluorescent frames (or consecutive frame numbers), and impute the length of this visit to be the time elapsed over *L*−1 frames, to obtain auxiliary quantities *f_k_* : time spent in fluorescent state (*k*th visit).

Up to discrete-approximation error, each *f_k_* represents the minimum of two exponential random variables with respective rates *r_D_* and *r_B_*, with likelihood contribution *p*(*f_k_*) = (*r_D_* + *r_B_*)exp{−(*r_D_* + *r_B_*)*f_k_*}.

Similarly, let *d_k_* denote the time elapsed over the *k*th interval between noncontiguous frames, taken to represent

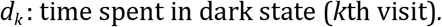

The likelihood contribution is

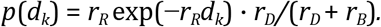

Finally, let *u* denote the time since the last blink (a period during which it is unknown whether the process has entered a dark or bleached state). The final contribution is

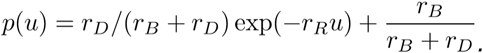

To finalise calculations, one must account for background noise (in the case *m* = 1). Such points are assumed to be uniform in spacetime. The complete marginal likelihood is:

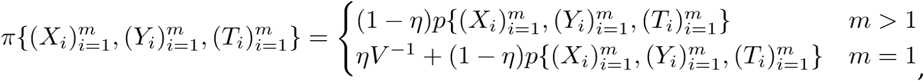

where *V* = *T*(*x*_0_ − *x*_1_)(*y*_1_ − *y*_0_), *T* is the length of the period of observation, and *η* is the background probability.

#### 2. Identifying and summarizing clusters

For both MBC and DTT clustering, an expected number of clusters, *N*, is first estimated, and a version of agglomerative hierarchical clustering (AHC) is then used to partition the dataset into *N* clusters. In AHC, each point is initially considered to be a distinct cluster. Using a user-specified metric and a linkage criterion, a stepwise greedy merging of the closest clusters is repeated until a partition with a predetermined number of clusters is obtained, or until no more clusters can be merged with a distance less than some specified number. The metric determines the distances between pairs of points, and the linkage criterion generalises these to a distance between clusters. Once the final partition has been identified, we merge each cluster down to its estimated centre, and the uncertainty of the centre is computed. In the following, we use the notation

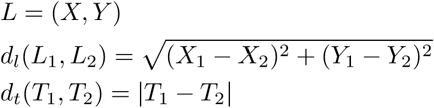

#### 3. MBC clustering

For MBC, the number of desired clusters, *N*, is an output of the rate-estimation step, and is thus decoupled from the clustering problem. For the AHC step, we use the family of metrics

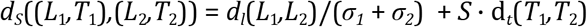

for *S* ≥ 0, which is simply the sum of the (scaled) Euclidean distance between the locations and times. For the linkage criterion we chose Ward’s Minimum Variance Method, as implemented via the Lance-Williams formula^17^, as it tends to find homogeneous clusters of spherical shape.

By varying *S*, we obtain a sequence of partitions, each slightly different but all chosen to have *N* clusters. The marginal likelihood is computed for each of the resulting partitions, and the most likely partition is selected.

#### 4. DTT clustering

DTT, or dark time thresholding, is a general idea in SMLM blinking correction literature, but implementation details are rarely discussed. The general principle is to merge locations that are close in space and time, with hard thresholds on the maximally allowed bridging distances in space and time. As a way to implement this idea in the AHC framework, we define the distance between 2 observations as

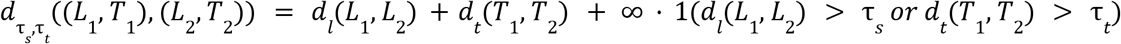

where ∞·0 = 0. Although not strictly a metric, this distance measure allows us to implement the dark time thresholding idea. We use the single-linkage criterion for cluster merging, which considers the distance between two clusters to be the smallest pairwise distance between them. Combined with our metric, this means that the clustering algorithm is allowed to merge points and clusters, so long as they can be combined via paths that do not violate the hard thresholds. Finally, a clustering is achieved by continuing to merge clusters until only infinite distances between clusters remain (no more legal merges can be made). For *τ_s_*, we used 4 times the mean localisation uncertainty. The temporal threshold, *τ_t_*, was determined as follows. First, the method of Annibale^4^ was used to determine *N*. Next, *τ_t_* was increased incrementally until the AHC algorithm produced a partition with *N* clusters, or as close to *N* as possible.

#### 5. Cluster centres and uncertainty

Let 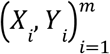 be the coordinates of an arbitrary cluster with centre *μ*. Once a particular clustering is given, it makes sense to treat the cluster centres as fixed parameters to be estimated. Thinking therefore of *μ* as fixed, the maximum likelihood estimator, 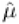, maximizes the likelihood of the cluster coordinates

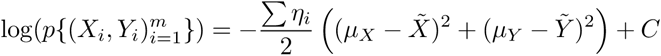

where *C* does not depend on *μ*, and it follows immediately that

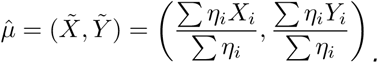

Using 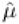, we can estimate the position of the molecule associated with a given cluster. As the coordinates of 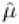 are independent, the covariance matrix of 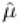 is given as

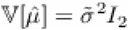

where *I*_2_ is the 2 × 2 identity matrix, and

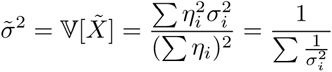

and the updated localisation uncertainty is then simply the associated standard deviation

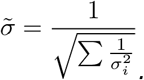

#### 6. Modifications to the rate-estimation procedures

The estimation of rates in the 4-state model is done using the methods of Jensen^8^. Using the notation of this work in the following, we here discuss an automatic way to select the set *U*, which should ideally cover large quantiles of the “survival time” distribution of a typical fluorophore (time from activation to permanent photobleaching). Although a suggestion for the selection of *U* is given in Jensen^8^, this requires that the model be fit twice, and depends on a standard assumption on the lifetimes of PALM fluorophores, and can thus lead to estimates of lowered quality in situations of atypical blinking.

To select an informative *U* in an automatic, data-driven way, we ideally wish to consider pairs of points from the same blinking cluster. As the probability of this event increases with spatial proximity, we consider the following statistic, which sums over pairs of nearest neighbors:

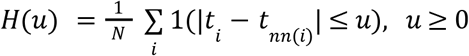

where *N* is the total number of observed datapoints, *t_i_* is the time (measured in seconds) associated with point *i*, and *nn*(*i*) is the index of the nearest neighbor (in space) to point *i*. Although the effect of background noise on this statistic can be quantified, it plays only a small effect in practice, and complicates the mathematical exposition, so we ignore it here. Computing the mean of *H*(*u*) can be done approximately by splitting the summation into 2 terms, where the first term corresponds to pairs (*i*, *nn*(*i*)) coming from the same blinking cluster, and the second term corresponds to pairs coming from different blinking clusters. Denote by *N*_s_ the set of index *i* ∈ {1, 2,.., *N*} such that point *nn*(*i*) belongs to the same blinking cluster as point *i*, and let *N*_d_ = {1, 2,.., *N*} − *N_s_* be the remaining index in {1, 2,.., *N*}. Then

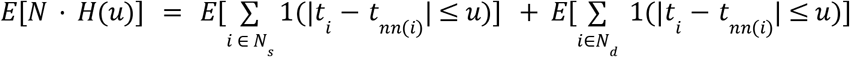

Let *X* be the set of protein locations, and denote by *G_x_* the number of points in the blinking cluster associated with the protein at location *x*. Further, let *x_i_* be the index of point *i* in the blinking cluster with centre *x*. Dealing with the first term, we condition on (*G_x_*, *X*) to obtain

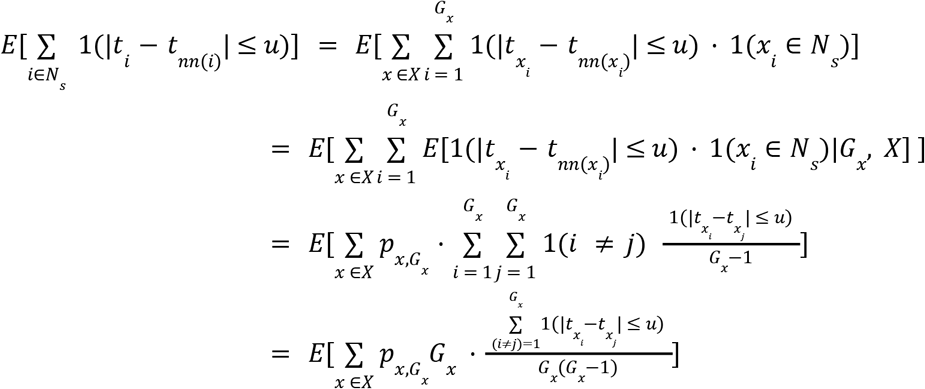

where by 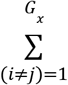 we denote the sum of all distinct pairs with index between1 and *G_x_*, and 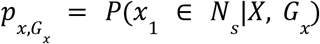. In going from the second to the third line we used that, given the event (1(*x_i_* ∈ *N_s_*) = 1) and *G_x_*, the timepoint 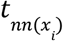 is randomly sampled among all *G_x_* − 1 timepoints in the blinking cluster at x different from 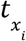, as the locations and times in a blinking cluster are independent conditional on *G_x_*, and further that the probability *P*(*x_i_* ∈ *N_s_*| *X*, *G_x_*) does not depend on *i*, because given (*X*, *G_x_*) the locations in a blinking cluster are identically distributed. Using a simple Taylor approximation, we thus obtain

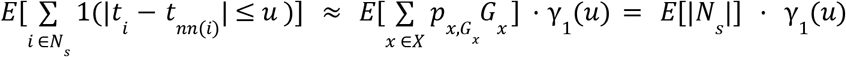

where |*N_s_*| denotes the number of elements in *N_s_*, and 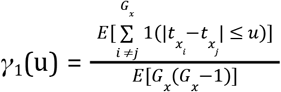 for arbitrary *x*. The identity 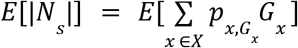 follows by setting *u* = ∞ in the above. Treating the second term in *E*[*N* · *H*(*u*)]in the same manner, we get

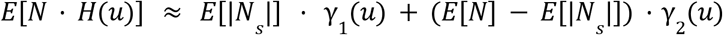

where γ_2_ (*u*), and its estimator ^^^γ_2_ (*u*), are as defined in Jensen^8^. Finally, using a second Taylor expansion we split up the LHS, so that

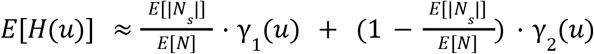

In the above, *u* ↦ γ_1_(*u*) is the distribution function of a non-negative random variable, and constitutes one part of the basis for rate parameter estimation in Jensen^8^. *U* should therefore be chosen to cover the time interval over which γ_1_ (*u*) moves between 0 and 1. Using the expression for *E*[*H*(*u*)] above, we do this in the following way. First, we compute the empirical distribution function *H*(*u*), and, setting *H*(*u*)in relation to its mean, we construct the loss-function

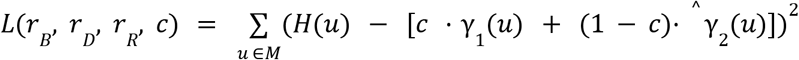

where the rates control the shape of γ_1_ (*u*), and *M* is the set consisting of the 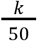-quantiles of the empirical *H*(*u*), for *k* ∈ {1, 2,…, 49}. The number of quantiles to consider can be chosen arbitrarily, with more typically leading to slightly better solutions, but slowing down optimization. In minimizing this loss-function, the rates are required to be positive, and *c* ∈ [0, 1]. The loss function is not informative enough to result in accurate rate estimates, but it does result in an estimate of γ_1_ (*u*), which we use to select an informative *U*. We do this by setting *U* to be the 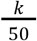-quantiles of the estimated γ_1_ (*u*), which ensures that the range of informative timepoints are adequately explored.

Again, more quantiles can be considered, if time permits.

#### Significance testing

The p-values reported in Supplementary Table 1 are based on a permutation test of the difference of means, using 10,000 simulations.

---

**Supplementary Table 1.** Summary of means and p-values for experimental data analysis of WT LAT and YF LAT at the immunological synapse.

|  | WT<br>Centre<br>mean | WT<br>Periphery<br>mean | YF Centre<br>mean | YF<br>Periphery<br>mean | WT<br>Centre vs<br>YF Centre<br>p-value | WT<br>Periphery<br>vs YF<br>Periphery<br>p-value | WT<br>Centre vs<br>WT<br>Periphery<br>p-value | YF Centre<br>vs YF<br>Periphery<br>p-value |
| --- | --- | --- | --- | --- | --- | --- | --- | --- |
| Number of<br>clusters | 10.0 | 16.5 | 4.76 | 4.25 | 0.006 | $< 10^{-5}$ | 0.043 | 0.701 |
| Number of<br>molecules<br>per cluster | 29.7 | 18.3 | 42.1 | 4.26 | 0.574 | 0.001 | 0.316 | 0.026 |
| Cluster<br>radius (nm) | 73.5 | 52.5 | 100 | 38.6 | 0.372 | 0.167 | 0.075 | 0.027 |
| Percentage<br>of<br>molecules<br>in clusters | 22.6 | 21.8 | 35.2 | 23.4 | 0.151 | 0.568 | 0.761 | 0.172 |
| Total<br>number of<br>molecules | 931 | 866 | 226 | 79.6 | $< 10^{-5}$ | $< 10^{-5}$ | 0.794 | 0.003 |
| Relative<br>density of<br>clustered vs<br>unclustered<br>molecules | 15.7 | 20.8 | 53.0 | 217 | $< 10^{-5}$ | $< 10^{-5}$ | 0.162 | 0.006 |

**Supplementary Figure 1:**
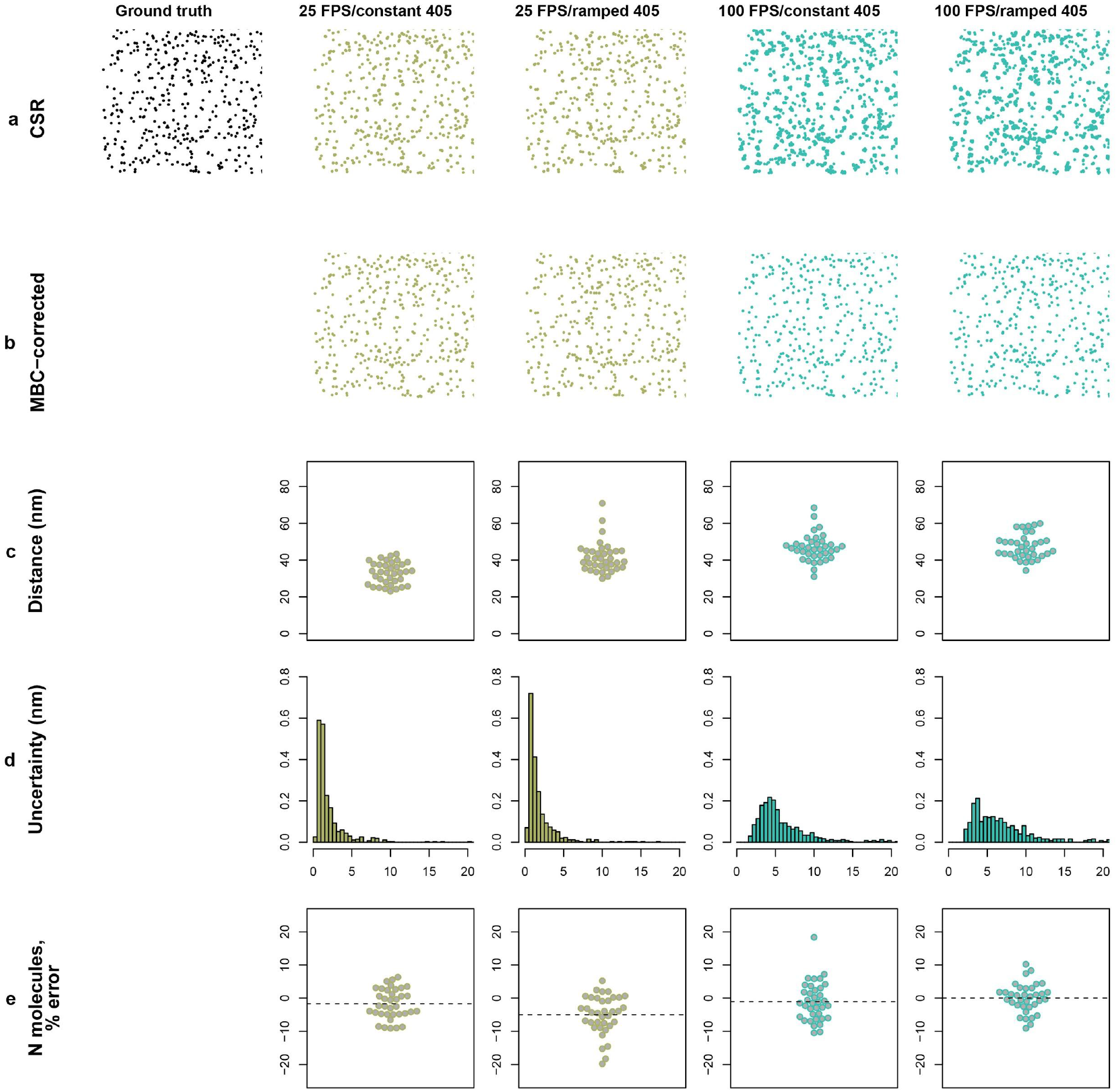
MBC Performance as a function of camera frame rate and the activating, 405 nm laser power. a) Example ground-truth and raw localisation maps for the different conditions. b) Example MBC-corrected maps. c) Wasserstein distances. c) Normalised histograms of localisation uncertainty. d) Percentage error in estimated number of ground-truth molecules (mean in dashed line).

**Supplementary Figure 2:**
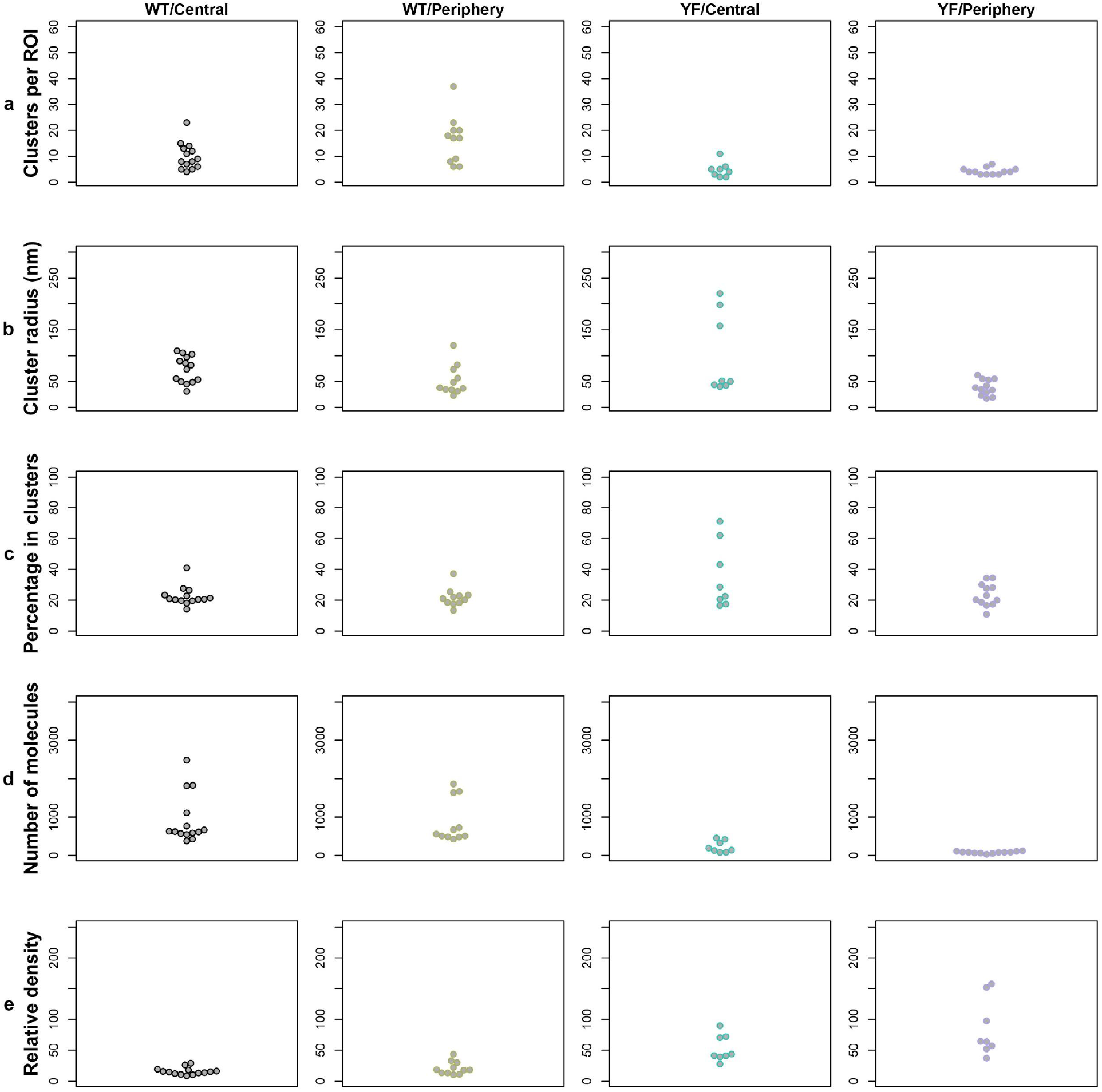
Additional statistics from the Bayesian cluster analysis of non-CSR LAT-mEos3.2 regions. a) Number of detected clusters, b) cluster radii (nm), c) percentage of molecules in clusters, d) number of molecules per ROI and e) relative density of molecules located in clusters as compared to the surrounding region.

