## Supplementary material for "Correction of multiple-blinking artefacts in photoactivated localisation microscopy": Code and example data: Dependencies.R

```
library(tidyverse)
library(magrittr)
library(wrapr)
library(spatstat)
library(multiway)
library(fastcluster)
library(dbscan)

source(file.path("Dependencies", "Estimation.R"))
source(file.path("Dependencies", "Gamma1.R"))
source(file.path("Dependencies", "Misc.R"))
source(file.path("Dependencies", "Postprocessing.R"))
source(file.path("Dependencies", "Scoring.R"))
source(file.path("Dependencies", "STBHClus.R"))
```
