## Supplementary material for "Correction of multiple-blinking artefacts in photoactivated localisation microscopy": Code and example data: Estimation.R

```
# Estimation functions ----
# --
# Fast computation of pair correlation function, based on the K function.
pcf.c = function(X,fast = T, correction = "iso", method = "c",...) {
  if(fast) {
    p = pcf.fv(Kest(X,correction = correction,...), method = method)
    p$iso = p$pcf
  } else {
    p = pcf(X,...)
  }
  return(p)
}

# PCF estimate with informative range of r-values.
trimmed_pcf = function(X, STDS, ...) {
  dnb = nndist(X); dnb = dnb[dnb > 0]
  
  rlow = mean(STDS)/4
  rmax = mean(STDS)*10
  rinc = (rmax-rlow)/5e2 # 500 equally spaced query points. Should be enough for realistic cases.
  
  pcf = pcf.c(unmark(X), r = seq(0,rmax, by = rinc), ...)
  range = which(pcf$r > rlow)
  
  return(list("PCF" = pcf, "Range" = range))
}

# Markcorrelation function based on markcorrelation integral.
Kmark.c = function(X,f, method = "c",...) {
  p = pcf.fv(Kmark(X,f, ...), method = method)
  p$iso = p$pcf
  return(p)
}

# Estimation of S_f for various f, given a pair-correlation function.
sumstat.get = function(X, f, r = NULL, plt = F, normalise = F, type = "K", prt = F) {
  zeroin = (0 %in% r | is.null(r)); if(!zeroin) {r = c(0,r)}
  if(is.null(r)) {r = pcf.c(X)$r}
  
  if(is.list(f)) {
    nf = length(f)
    for(v in 1:length(f)) {attributes(f[[v]])$name = paste("- Computing summary statistic ", v, " of ", nf, ".", sep = "")}
    o = sapply(f, function(L) sumstat.get(X, L, r, normalise = normalise, type = type, prt = prt))
  } else {
    if(prt){overprint(attributes(f)$name)}
    if(type == "M") {
      mkuv = markcorr(X, f = f, r = r, correction = "iso", normalise = normalise)
      mkuv = mkuv$iso
    } else {
      mkuv = Kmark.c(X, f = f, r = r, correction = "iso", normalise = normalise)
      mkuv = mkuv$iso
    }
    
    return(mkuv)
  }
  catprint("")
  out = t(unlist(o))
  if(!zeroin) {out = out[,-1]; r = r[-1]}
  if(plt) {if(is.null(dim(out))) {plot(out~r, type = "l")} else {matplot(y = t(out), x = r, type = "l")}}
  return(out)
}

# Generates a list of functions on the form 1(|x-y| <= u)
funlistgen.u = function(u) {
  ffu = function(v) return(function(m1,m2) abs(m1-m2) <=  v)
  lf = list(); for(uv in u) {lf = append(lf, do.call("ffu", list(uv) ))}
  return(lf)
}
# Generates a list of functions on the form 1(x <= u)
funlistgen.q = function(q) {
  ffq = function(v) return(function(m1,m2) m1 <=  v)
  lf = list(); for(qv in q) {lf = append(lf, do.call("ffq", list(qv) ))}
  return(lf)
}

U.G.Get = function(X, g2s, framerate, nstep = 50) {
  d = X %$% data.frame(x,y,marks)

  o = dbscan::kNN(d[,1:2], k = 1)
  st = c()
  n = nrow(d)
  for(i in 1:n) {
    m1 = d[i,3]
    m2 = d[o$id[i],3]
    
    st = c(st, abs(m1-m2))
  }
  ee = ecdf(st)
  us = approxfun(ee(st), st, rule = 2)(seq(0,0.95,length.out = nstep))
  
  us = unique(round(us*framerate))/framerate
  obs = ee(us)
  
  g2  = g2s$g2(us)
  g22 = cbind(g2)
  
  plot(obs~us, type = "l", ylim = c(0,1))
  points(g2~us, type = "l", col = "red")
  
  f = function(par, geta = F) {
    g1 = dist1approx(us, par, 1/framerate)
    fw = cbind(g1,g22)
    
    a = multiway::fnnls(crossprod(fw), crossprod(fw, obs))
    a = a/sum(a)
    if(geta) {
      return(a)
    }
    o = mean((obs-fw%*%a)^2)
    return(o)
  }
  
  o = boptim.try(c(3,2), f, n = 20, plt = F)
  pr = abs(o$par)

  g1f = function(u) dist1approx(u, pr, 1/framerate)
  
  up = 0
  a = 0
  while(a < 0.99) {
    a = g1f(up)
    up = up+1/framerate
  }
  
  gxd = seq(0,up, length.out = 250)[-1]
  gxd = approxfun(g1f(gxd), gxd, rule = 2)(seq(min(obs),0.99,length.out = nstep+1))[-1]
  gxd = unique(round(gxd*framerate))/framerate
  gxd = gxd[gxd >= 1/framerate]

  abline(v = gxd, lty = 2, col = "gray")
  
  points(g1f(us)~us, type = "l", col = "green")

  a = f(o$par, geta = T)
  points(cbind(g1f(us),g22)%*%a~us, type = "l")
  
  return(list(par = pr, g1 = g1f(gxd), us = gxd, weights = a, obs = ee(gxd), g2 = g2s$g2(gxd)))
}

g2.estim = function(M,alpha, R = max(M), plt = F, nsim = 1e5) {
  MM = sort(M)
  uM = unique(MM)
  oecdf = ecdf(MM)(uM)
  
  uM  = c(0,uM); oecdf = c(0,oecdf)
  
  necdf = uM/R
  g2ecdf = (oecdf-(1-alpha)*necdf)/alpha
  
  fq.o = approxfun(y = uM, x = oecdf)
  fq = approxfun(y = uM, x = g2ecdf)
  
  ss.g2o = abs(fq.o(runif(nsim,0,1))-fq.o(runif(nsim,0,1)))
  og.g2o = sapply(uM, function(u) mean(ss.g2o <= u))
  
  ss.g2 = abs(fq(runif(nsim,0,1))-fq(runif(nsim,0,1)))
  og.g2 = sapply(uM, function(u) mean(ss.g2 <= u))
  
  og.ne = function(x) 1-(1-x/R)^2
  
  ss.mix = abs(fq(runif(nsim,0,1))-runif(nsim, 0, R))
  og.mix = sapply(uM, function(u) mean(ss.mix <= u))
  
  fo.g2o = approxfun(x = uM, y = og.g2o, rule = 2)
  fo.g2  = approxfun(x = uM, y = og.g2, rule = 2)
  fo.ne  = og.ne
  fo.mix = approxfun(x = uM, y = og.mix, rule = 2)
  
  if(plt) {
    foo = abs(sample(M,1e5, T)-sample(M,1e5, T))
    s0 = sapply(uM, function(u) mean(foo <= u))
    plot(fo(uM)~uM, type = "l", col = "green", lwd = 1.5, xlab = "u", ylab = "Fu")
    points(s0~uM, type = "l", col = "red", lwd = 1.5)
    legend(x = "bottomright", legend = c("gamma2", "gamma2o"), col = c("green", "red"), lty = 1)
  }
  return(list(g2o = fo.g2o, g2 = fo.g2, g2e = fo.ne, g2ez = fo.mix)) 
}

sigma.bias.estim = function(summat, sigma, r, r.range, alpha, into, sseq = seq(0.25, 2.75, by = 0.25/10)) {
  bst.s = 1
  bst.er = Inf
  for(sc in sseq) {
    At = get.A(r[r.range], sigma*sc, 1e3)
    W = apply(summat, 1, function(s) funsplit(s, At*alpha/into, 0, fixbc = T))
    er = mean((summat-W%*%t(At)*alpha/into)^2)
    if(er < bst.er) {
      bst.er = er
      bst.s = sc
    }
  }
  return(bst.s)
}

correct.inrate = function(r0,b) {
  f = function(l) (exp(l*b)-l*b-1)/(l*(exp(l*b)-1))-1/r0
  xd = uniroot(f, c(r0/10, r0))
  return(xd$root)
}

noise.estim = function(X,pcf, r.range, ur = min(X$marks)+max(X$marks), ...) {
  noisesummary = sumstat.get(X, list(function(m1,m2) m1+m2), r = pcf$r, normalise = F)
  stat = noisesummary[r.range]
  gg = pcf$iso[r.range]
  r = pcf$r[r.range]
  
  ao = mean(X$marks)*2
  opt1 = funsplit(stat, A = gg-1, ao, fixbc = T)
  
  a = (ao-ur)/(opt1-ur)

  out = clamp(a, 0, 1)

  return(c(out,ao))
}

optf = function(par, u, O, g2, fr) {
  par = abs(par)
  g1 = dist1approx(u, par, 1/fr)
  nc = moment.approx(par, 1/fr)[3]
  guess = (g1-g2)*nc
  
  loss = mean((O-guess)^2)
  return(loss)
}

optf.G = function(par, u, O, ming1, g2, fr) {
  par = abs(par)
  g1 = dist1approx(u, par, 1/fr)
  nc = moment.approx(par, 1/fr)[3]
  guess = (g1-g2)*nc
  
  loss  = mean((O-guess)^2)+1
  loss2 = sum((g1 < ming1)*(ming1-g1)) # Ensuring gamma1 lies on the correct side of H(u)
  return(loss+loss2)
}

# Estimate all.
# 
estimate.model = function(X, framerate, sig = NA, alpha = 1, ndark = 1, cut = NA, correct.sigma.bias = F, nopt = 50, nu = 50, ...) {
  par(mfrow = c(3,2), mai = rep(0.4,4))
  if(!all(is.integer(X$marks))) {catprint("Please provide point pattern with frame-marks."); stop("Non-integer marks detected.")}
  
  hist(X$marks/framerate, breaks = min(round(X$n/10), 100))
  if(!any(is.na(cut))) {
    X = X %$% subset(., between(.$marks, cut[1], cut[2]))
    X$marks = X$marks-cut[1]
    catprint("Data was subset to cut range.")
    abline(v = cut/framerate, lty = 2, col = "red")
  }
  X$marks = X$marks/framerate
  
  breakl(); catprint("Estimating gamma1 and obtaining U.")
  g2s = g2.estim(X$marks, alpha)
  
  r1st = U.G.Get(X, g2s, framerate, nu)
  u = r1st$us
  
  g2u = g2s$g2(u)
  
  breakl(); catprint("Estimating PCF.")
  into = intensity(X)
  pcf.l = trimmed_pcf(X, sig)
  
  pcf = pcf.l$PCF$pcf
  r = pcf.l$PCF$r
  r.range = pcf.l$Range
  
  plot(pcf[r.range]~r[r.range], type = "l", main = "PCF: trimmed")
  
  breakl(); catprint("Estimating alpha.")
  if(is.na(alpha)) {
    foo = noise.estim(X, pcf.l$PCF, r.range, plt = T)
    alpha = foo[1]
    m0 = foo[2]
  } else {
    m0 = mean(X$marks)*2
  }
  estprint(alpha, "alpha")
  
  breakl(); catprint("Estimating summary statistics.")
  
  g2o = sapply(u, function(us) mean(replicate(100, mean(abs(sample(X$marks)-sample(X$marks)) <= us))) )
  
  summat = sumstat.get(X, funlistgen.u(u), r[r.range], plt = T, prt = T, normalise = F)
  
  for(i in 1:nrow(summat)) {
    summat[i,] = summat[i,]-(pcf[r.range]-1)*g2u[i]-g2o[i]
  }
  matplot(t(summat), x = r[r.range], type = "l")
  
  bst.s = 1
  if(correct.sigma.bias) {
    breakl(); catprint("Estimating sigma bias.")
    
    bst.s = sigma.bias.estim(summat, sig, r, r.range, alpha, into)
    A = get.A(r[r.range], sig*bst.s, 1e4)
    catprint(c("Sigma bias scaling:", bst.s ))
  } else {
    A = get.A(r[r.range], sig, 1e4)
  }
  
  breakl(); catprint("Estimating rate-parameters.")
  
  W = apply(summat, 1, function(s) funsplit(s, A*alpha/into, 0, fixbc = T))
  
  f = function(par) optf.G(par, u, W, r1st$obs, g2u, framerate)
  
  ratepar = boptim.try(c(3, 4), f, nopt, mean = clamp(r1st$obs, 0, framerate*2))
  
  ratepar = abs(ratepar$par)
  parinfo = pars.to.terms(ratepar)
  
  lr = 1/((mean(X$marks)-(1-alpha)*sum(range(X$marks))/2)/alpha-gam2d.AB(ratepar, 1/framerate))
  #lrc = correct.inrate(lr, max(X$marks))
  lrc = 0
  
  ratepar = c(ratepar, lr, lrc)
  stt = paste(c("r-FD","r-DF"), sort(rep(1:ndark,2)), sep = "")
  names(ratepar) = c("r-FBL", stt, "r-IFL", "r-IFL-cor")
  
  estprint(ratepar, names(ratepar), 4)
  
  breakl(); catprint("Estimation complete!")
  return(list(par = ratepar, noise = alpha, sig.scale = bst.s))
}

# estimate.model.nosig = function(X, framerate, rmax, alpha = 1, ndark = 1, cut = NA, nopt = 50, nu = 50, ...) {
#   par(mfrow = c(3,2), mai = rep(0.4,4))
#   if(!all(is.integer(X$marks))) {catprint("Please provide point pattern with frame-marks."); stop("Non-integer marks detected.")}
# 
#   hist(X$marks/framerate, breaks = min(round(X$n/10), 100))
#   if(!any(is.na(cut))) {
#     X = X %$% subset(., between(.$marks, cut[1], cut[2]))
#     X$marks = X$marks-cut[1]
#     catprint("Data was subset to cut range.")
#     abline(v = cut/framerate, lty = 2, col = "red")
#   }
#   X$marks = X$marks/framerate
# 
#   breakl(); catprint("Estimating gamma1 and obtaining U.")
#   g2s = g2.estim(X$marks, alpha)
#   r1st = U.G.Get(X, g2s, framerate, nu)
# 
#   u = r1st$us
#   g2u = g2s$g2(u)
#   g11 = r1st$g1
# 
#   breakl(); catprint("Estimating g1nc.")
#   into = intensity(X)
# 
#   g2o = sapply(u, function(us) mean(replicate(100, mean(abs(sample(X$marks)-sample(X$marks)) <= us))) )
# 
#   W2 = Kmark(X, function(m1,m2) abs(m1-m2)/fr <= Inf, r = c(0,rmax), normalise = F, correction = "iso")$iso[2]*into/alpha
#   Wf = function(i) Kmark(X, function(m1,m2) abs(m1-m2) <= u[i], r = c(0,rmax), normalise = F, correction = "iso")$iso[2]*into/alpha
#   W.un = sapply(1:length(u), Wf)
#   W = W.un-g2u*W2-g2o
# 
#   plot(W.un~u, type = "b")
#   plot(W~u, type = "b")
# 
#   breakl(); catprint("Estimating rate-parameters.")
# 
#   f = function(par) optf.G(par, u, W, g2u, framerate)
# 
#   ratepar = boptim.try(c(1+2*ndark, 4), f, nopt, mean = abs(r1st$par))
# 
#   fitqual = exp(ratepar$val)
#   ratepar = abs(ratepar$par)
#   parinfo = pars.to.terms(ratepar)
# 
#   lr = 1/((mean(X$marks)-(1-alpha)*sum(range(X$marks))/2)/alpha-gam2d.AB(ratepar, 1/framerate))
#   #lrc = correct.inrate(lr, max(X$marks))
#   lrc = 0
# 
#   ratepar = c(ratepar, lr, lrc)
#   stt = paste(c("r-FD","r-DF"), sort(rep(1:ndark,2)), sep = "")
#   names(ratepar) = c("r-FBL", stt, "r-IFL", "r-IFL-cor")
# 
#   estprint(ratepar, names(ratepar), 4)
# 
#   breakl(); catprint("Estimation complete!")
#   return(list(par = ratepar, noise = alpha))
# }
# # 
# # 
# # 
# # 
# # 
# # 
# # 
# # 
# # 
# # 
```
