## Supplementary material for "Correction of multiple-blinking artefacts in photoactivated localisation microscopy": Code and example data: Gamma1.R

```
phase_type_interval = function(par, alpha = 0.025) {
  lb = par[1]
  ld = par[2]
  lr = par[3]
  m = (lr+ld)/((lb+ld)*lr-ld*lr)
  v1 = 1/(lb^2*lr)*(ld+lr+(lb*ld+ld^2+ld*lr)/lr)
  v = 2*v1-m^2

  r = m+sqrt(v/alpha)
  return(r)
}

mgf = function(v, lamb) {
  lamb/(lamb-v)
}

mgn = function(v, p) {
  p*exp(v)/(1-(1-p)*exp(v))
}

gam2d.AB = function(par, delt) {
  pt = pars.to.terms(par)
  GM = pt$mf/delt+1/2
  A = 1/GM*(pt$mf2/(2*delt)+ pt$mf+3*delt/8)
  B = 1/(pt$mninf*GM)*( pt$mf/delt+1/2 )*(1/2*(pt$mninf2-pt$mninf)*(pt$mf+pt$mr)+pt$mninf*delt*1/2)
  return(A+B)
}

gam1.asymp.x = function(v,mgff,mgfr,mninf,mninf2,mf,mf2, spec) {
  mgFR = mgfr(-v)*mgff(-v)
  enum = mninf*(mgff(-v)+v*mf-1)+((mgff(-v)-1)/(mgFR-1))^2*mgfr(-v)*(spec(v)-mgFR*mninf+mninf-1  )
  denom = v^2*(mninf*mf2 + mf^2*(mninf2-mninf))
  o = 2*(enum)/denom
  return(o)
}

gam1.asymp = function(v, par) {
  terms = pars.to.terms(abs(par))
  o = gam1.asymp.x(v, terms$mgff, terms$mgfr, terms$mninf, terms$mninf2, terms$mf, terms$mf2, terms$spec)
  return(o)
}

gam1.asymp.d1 = function(v,par) {
  par = abs(par)
  lB = par[1]
  lF = par[2]
  lR = par[3]
  lO = lF+lB
  out = (lB*(lR+v))/(lB * lR+(lB+lF+lR)*v+v^2)
  
  return(out)
}

gam1approx.x = function(v,mgff,mgfr,mninf,mninf2,mf,mf2, spec, delt) {
  eps = 1; teps = 1
  mgFR = mgfr(-v)*mgff(-v)
  a = 2*mninf*(exp(delt*v)*(mgff(-v)*exp(-delt*v*eps)-1) + (exp(delt*v)-1)*(mf/delt+eps))*1/(1-exp(delt*v))^2
  b = mgfr(-v)*(spec(v)-mgFR*mninf+mninf-1)/(1-mgFR)^2
  c = 2*(exp(delt*v*(1+teps)))*(1-mgff(-v)*exp(-delt*v*eps))^2*1/(1-exp(delt*v))^2
  d = mninf2*(mf/delt+eps)^2+mninf*((mf2-mf^2)/delt^2-mf/delt-eps)
  return((a+b*c)/d)
}

gam1approx = function(v, par, delt) {
  terms = pars.to.terms(abs(par))
  o = gam1approx.x(v, terms$mgff,terms$mgfr,terms$mninf,terms$mninf2,terms$mf,terms$mf2, terms$spec, delt)
  o[is.nan(o)] = 1
  return(o)
}

# Old version with fixed quality settings. 
# dist1approx = function(u, par, delt, eps = 1e-2, k = 0:1e3){ #, eps = 1e-2, k = 0:(1e3)) {
#   O = gam1approx(-1i*(k+1/2)*eps, par, delt)
#   fx = function(x) 1/2-sum(Im(O*exp(-1i*(k+1/2)*eps*x))/(pi*(k+1/2)))
#   
#   a = sapply(u, fx)
#   return(a)
# }

# Automatically selects inversion quality settings.
dist1approx = function(u, par, delt){
  if(all(u == 0)) {return(rep(0,length(u)))}
  # vvv Picking quality settings to match parameters
  mu = max(u)
  
  gm = phase_type_interval(abs(par))
  
  m = max(c(mu,gm-mu))
  
  eps = 2*pi/(m+1)
  km = min(c(round(1e2/(eps)), 2e3))
  
  k = 0:km
  # Picking quality settings to match parameters
  
  # ^^^ Performing the inversion
  O = gam1approx(-1i*(k+1/2)*eps, par, delt)
  fx = function(x) 1/2-sum(Im(O*exp(-1i*(k+1/2)*eps*x))/(pi*(k+1/2)))

  a = sapply(u, fx)
  return(a)
}

# Laplace-transform based inversion. Don't use this.
# dist1approx_laplace = function(u, par, delt) {
#   LT = function(v) (gam1approx(v, par, delt)/v)
#   
#   umi = max(c(min(u)%/%delt, 1))
#   uma = max(u)%/%delt+1
#   
#   o = pracma::invlap(LT, umi*delt, uma*delt, uma-umi+1)$y
#   o = c(0, o)
#   
#   out = o[pmax(u%/%delt, 1)-umi+2]
#   
#   return(out)
# }

dist1asymp = function(u, par, eps = 1e-2, k = 0:(1e3)) {
  O = gam1.asymp(-1i*(k+1/2)*eps, par)
  fx = function(x) 1/2-sum(Im(O*exp(-1i*(k+1/2)*eps*x))/(pi*(k+1/2)))
  
  a = sapply(u, fx)
  return(a)
}

meanapprox = function(par, delt, s = 0.01) {
  return(-diff(gam1approx(c(-s,s), par, delt))/(2*s))
}

moment.approx = function(par, delt) {
  pt = pars.to.terms(par)
  
  nsub = 1000
  mur1 = integrate(function(x) (1-x)*pt$pdfr(x*delt)*delt, 0, 1, subdivisions = nsub)$value
  mur2 = integrate(function(x) (1-x)^2*pt$pdfr(x*delt)*delt, 0, 1, subdivisions = nsub)$value
  
  m1 = pt$mninf*(pt$mf/delt+1)-(pt$mninf-1)*mur1
  
  m2 = 
    pt$mninf2*(pt$mf/delt+1)^2+pt$mninf*(pt$mf2-pt$mf^2)/delt^2+
    (pt$mninf2+1-2*pt$mninf)*mur1^2+(pt$mninf-1)*(mur2-mur1^2)-
    2*(pt$mf/delt+1)*mur1*(pt$mninf2-pt$mninf)
  
  ov = c(m1,m2,m2/m1-1)
  names(ov) = c("M1", "M2", "nc")
  return(ov)
}

pars.to.terms = function(par) {
  par = abs(par)
  nd = (length(par)-1)/2
  
  rFB = par[1]
  par = par[-1]
  
  out.rates  = par[seq(1,nd*2, by = 2)]
  back.rates = par[seq(2, nd*2, by = 2)]
  
  rOut = sum(out.rates)+rFB
  p = rFB/rOut
  
  mix = out.rates/sum(out.rates)
  
  mninf = 1/p
  mninf2 = (1-p)/p^2+1/p^2
  
  mf = 1/rOut
  mf2 = 2/rOut^2
  
  mr = 0; for(i in 1:nd) {mr = mr+mix[i]*1/back.rates[i]}
  
  mgff = function(v) mgf(v,rOut) 
  mgfr = function(v) {
    o = 0
    for(i in 1:nd) {
      o = o+mix[i]*mgf(v,back.rates[i])
    }
    return(o)
  }
  CDFr = function(v) {
    o = 0
    for(i in 1:nd) {
      o = o+mix[i]*pexp(v,back.rates[i])
    }
    return(o)
  }
  PDFr = function(v) {
    o = 0
    for(i in 1:nd) {
      o = o+mix[i]*dexp(v,back.rates[i])
    }
    return(o)
  }
  spec = function(v) mgn(log(mgff(-v)*mgfr(-v)),p)
  
  return(list(rOut = rOut,mf = mf, mf2 = mf2, mr = mr, mninf = mninf, mninf2 = mninf2, mgff = mgff, mgfr = mgfr, pdfr = PDFr, cdfr = CDFr, spec = spec, p = p, mix = mix, ndark = nd, out.rates = out.rates, back.rates = back.rates))
}

# gam1deltapprox = function(v, par, l, delt) { # gamma1 after addition of exponential noise with parameter lambda = l
#   g1 = gam1approx(v, par, delt)
#   g1l = rep(gam1approx(l, par, delt), length(v))
# 
#   out = (g1-g1l*v/l)/((1+v/l)*(1-v/l))
# 
#   if(l %in% v) {
#     wb = which(v == l)
#     oof = diff(gam1approx(c(l-0.01,l+0.01), par, delt, eps))/0.02
#     out[wb] = 1/2*(g1[wb]-l*oof)
#   }
#   return(out)
# }
# 
# dist1deltapprox = function(u, par, delt, l, eps = 1e-2, k = 0:(1e3)) {
#   if(l == 0) {return(rep(0, length(u)))}
#   O = gam1deltapprox(-1i*(k+1/2)*eps, par, l, delt)
#   fx = function(x) 1/2-sum(Im(O*exp(-1i*(k+1/2)*eps*x))/(pi*(k+1/2)))
#   
#   a = sapply(u, fx)
#   return(a)
# }
```
