## Supplementary material for "Correction of multiple-blinking artefacts in photoactivated localisation microscopy": Code and example data: Misc.R

```
# Helper functions ----
# --

# Nice printing.
overprint = function(s) {
  cat("\r",s)
}

catprint = function(s) {
  cat(s, "\n")
}

estprint = function(e,n,d = 3) {
  e = round(e, d)
  s = paste(n, "<-", e, sep = " ")
  for(si in s) {catprint(si)}
}

breakl = function() {
  catprint("______________________________________")
}

# Write line to csv file
write.to.file = function(l,file, main = NULL) {
  if(!file.exists(file)) {
    file.create(file)
    if(!is.null(main)) {
      write.table(main,file = file, append = T, col.names = F, row.names = F, quote = F)
    }
  } else {
    write.table(l,file = file, append = T, col.names = F, row.names = F, quote = F)
  }
}

# Multi-start optimization with performance graph.
boptim.try = function(starter.v, fun, n = 1e3, plt = T, silent = F, prt = F, use.sof = T, reset = 100, mean = 0, ...) { 
  if(length(starter.v) == 2) {starter = function() rnorm(starter.v[1],mean, starter.v[2]) } else {starter = starter.v}
  b = Inf
  avs = c()
  sof = rep(0,length(starter()))
  for(i in 1:n) {
    copt = tryCatch(optim(sof+starter(), fun, ...), error = function(e) "ER")
    if(copt[1] == "ER") {
      print(copt)
      n = n+1
      sof = rep(0, length(starter()))
      if(!silent) {
        print(paste("Function returned an error on iteration ",i, sep = ""))
      }
    } else {
      cval = copt$value
      avs[i] = cval
      if(cval < b) {
        params = copt$par
        b = cval
        if(prt) {print(c(round(params,3), log(b) ))}
        if(use.sof) {sof = params; if(any(sof > reset)) {sof = rep(reset, length(params))}}
      }
    }
    
  }
  if(plt) {
    hist(log(avs), breaks = round(min(c(n,100))))
    abline(v = log(b), col = "red", lty = 2, lwd = 2)
  }
  return(list(par = params, val = b))
}

boptim = function(starter, fun, n = 1e3, plt = T, ...) { 
  b = Inf
  avs = numeric(n)
  for(i in 1:n) {
    copt = optim(starter(), fun, ...)
    cval = copt$value
    avs[i] = cval
    if(cval < b) {
      params = copt$par
      b = cval
    }
  }
  if(plt) {
    hist(log(avs), breaks = min(c(n/3,100)))
    abline(v = log(b), col = "red", lty = 2, lwd = 2)
  }
  return(list(par = params, val = b))
}

# Clamps a value
clamp = function(v, l, u) {
  pmin(pmax(v, l), u)
}

# Splits a function into an affine combination of 2 given functions
funsplit = function(O,A,B, fixbc = F) {
  if(!fixbc) {
    M = cbind(A,B,1)
    S = t(M)%*%M
    out = solve(S)%*%t(M)%*%O; out = as.numeric(out)
    names(out) = letters[1:3]
  } else {
    M = cbind(A)
    S = t(M)%*%M
    out = solve(S)%*%t(M)%*%(O-B); out = as.numeric(out)
    names(out) = "a"
    
  }
  return(out)
}
funsplitx = function(O,A,B,C = NULL) {
  if(!is.null(C)) {
    M = cbind(A,B,C)
    S = t(M)%*%M
    out = solve(S)%*%t(M)%*%O; out = as.numeric(out)
    names(out) = letters[1:3]
    return(out)
  } else {
    M = cbind(A,B)
    S = t(M)%*%M
    out = solve(S)%*%t(M)%*%O; out = as.numeric(out)
    names(out) = letters[1:2]
    return(out)
  }
}

# Cluster term obtained via simulation from collection of uncertainties.
AN = function(r,s1,s2) {
  con = 2*pi*s1*s2*sqrt(1/s1^2+1/s2^2)*sqrt(s1^2+s2^2)
  rter = exp(-r^2/(2*(s1^2+s2^2)))
  return(1/con*rter)
}
get.A = function(r, sigdst, nsim = 1e3) {
  fx = function() {sv = sample(sigdst,2,T); o = AN(r, sv[1],sv[2]); return(o)}
  m = replicate(nsim, fx(), simplify = T)
  o = rowMeans(m)
  return(o)
}
APS = function(r,sigs,ns = 1e3) {
  s1 = sample(sigs, ns, T)
  s2 = sample(sigs, ns, T)
  
  S = cbind(s1,s2)
  
  o = apply(S, 1, function(v) AN(r,v[1],v[2]))
  return(rowMeans(o))
}
good_sample = function(v,size,replace,m) {
  if(length(v) == 0) {return(NA)}
  if(length(v) == 1) {return(v)}
  
  return(sample(v,size, replace))
}
splitf.A = function(A, r, B, sm) {
  o = t(apply( sm,1, function(k) funsplitx(k, A, B , 1) ))
  return(o)
}

splitf.A.noint = function(A, r, B, sm) {
  o = t(apply( sm,1, function(k) funsplitx(k, A, B) ))
  return(o)
}
# Data postprocessing / tools ----
# ---
# Marked for deletion!

# duplicate_remover = function(X,r) {
#   xm = sort(unique(X$marks))
#   xdf = X %$% data.frame(x = x, y = y, mar = marks)
#   out = data.frame()
#   
#   for(m in xm) {
#     wrp = xdf %>% filter(mar == m)
#     if(nrow(wrp) == 1) {
#       out = rbind(out, wrp)
#       next
#     }
#     
#     dt = dbscan::dbscan(wrp[,1:2], r, 1)
#     wrpn = wrp %>% group_by(dt$cluster) %>% sample_n(., 1) %>% ungroup() %>% select(-'dt$cluster')
#     
#     out = rbind(out, wrpn)
#     if(nrow(wrpn) < nrow(wrp)) {
#       print(paste("Eliminated merging on frame: ", m, sep = ""))
#     }
#     
#   }
#   out = out %$% ppp(x,y,marks = mar, window = X$window)
#   return(out)
# }
# 
# data_grouper = function(X,sig,q = 0.99) {
#   threshold = sig*sqrt(qchisq(q,2))
#   print(paste("Grouping distance threshold: ", round(threshold,3), sep = ""))
#   
#   XO = X %$% data.frame(x = x, y = y, m = marks)
#   
#   xm = sort(unique(XO$m))
#   out = data.frame()
#   
#   xmq = quantile(xm, seq(0,1,by = 0.1))[-c(1,10)]
#   for(mm in xm) {
#     if(mm %in% round(xmq,0) ) {print(paste("Processing frame:", mm, sep = ""))}
#     CCurr = XO %>% filter(m == mm)
#     CPrev = XO %>% filter(m == mm-1)
#     
#     if(nrow(CPrev) == 0) {
#       out = rbind(out, CCurr)
#       next
#     }
#     
#     OM = dbscan::frNN(CPrev[,1:2], threshold, CCurr[,1:2])
#     
#     wk = which(unlist(lapply(OM$dist, length)) == 0)
#     out = rbind(out, CCurr[wk,])
#   }
#   o = out %$% ppp(x = x, y = y, marks = m, window = X$window)
#   return(o)
# }
```
