## Supplementary material for "Correction of multiple-blinking artefacts in photoactivated localisation microscopy": Code and example data: PostProcessing.R

```
regionsplit = function(df,nsub = 5) { # Convenience function for splitting large ROIS into smaller blocks.
  xr = range(df[,1])
  yr = range(df[,2])
  
  spltx = seq(xr[1], xr[2], length.out = nsub+2)#[-c(1,nsub)]
  splty = seq(yr[1], yr[2], length.out = nsub+2)#[-c(1,nsub)]
  
  xbin = cut(df[,1], spltx)
  ybin = cut(df[,2], splty)
  
  obin = interaction(xbin,ybin)
  obin = as.numeric(obin)
  df$bin = obin
  return(split.data.frame(df, as.factor(df$bin)))
}

# Cluster and score, optimizing over the hyper-parameter.
# Now weights spatial distances by uncertainty!
STBHClus_optimize = function(df, framerate, ord.params, eint, t.thres = seq(0.005,0.08,length.out = 30), method = "ward.D2", plt = F) {
  best = -Inf
  clus = NA
  best.ind = 0
  uncert = df[,4]
  
  N = nrow(df)
  n.blink = moment.approx(ord.params[c(4,2,3)], 1/framerate)[1]
  
  EN =  diff(range(df[,1]))*diff(range(df[,2]))*eint
  
  n.expected.protein = round((N-EN)/n.blink)
  
  npar = length(t.thres)
  
  noise = EN/(n.expected.protein+EN)
  
  Nclus = round(n.expected.protein+EN)
  
  ord.params = clamp(ord.params, 0, 5*framerate) # fixing divergent parameters.
  s = 1/as.dist(outer(uncert, uncert, "+"))
  D1 = dist(df[,1:2])*s
  D2 = dist(df[,3])
  
  df[,3] = as.integer(df[,3])
  v.scores = c()
  for(j in 1:npar) {
    dts = t.thres[j]
    DUse = (D1)+(D2)*dts
    
    m = cluster_and_score(df[,1:3], STBHClus_N, score_STC, cluster.args = list(N = Nclus, D = DUse, method = method), 
                          score.args = list(sds = uncert, pars = ord.params, noise = noise, framerate = framerate))
    
    v.scores = c(v.scores, m$score)
    if(m$score > best) {
      best = m$score
      best.dt = dts
      #if(prt){print("- - - - -")}
      clus = m$clustering
    }
  }
  if(best.dt == last(dts) | best.dt == dts[1]) {print("Warning: best solution is an edge case. Consider a wider t.thres.")}
  if(plt) {
    plot(v.scores~ t.thres, type = "l", lwd = 1.5, col = "red")
    abline(v = best.dt, h = best, lty = 2)
  }
  merge = merge.clusters(df[,1:3], uncert, clus, keep.mix = T); merge = merge[,-6]
  return(list(merged = merge, clus = clus, loglik = best))
}

# Optimally summarize found clusters into singular localizations with new uncertainties.
merge.clusters = function(pts, sds, labels, keep.mix = T) {
  df = data.frame(x = pts[,1], y = pts[,2], time = pts[,3], labels = labels, w = 1/sds^2)
  
  dfgroup = df %>% group_by(labels) %>% summarise(x = sum(w*x)/(sum(w)),  y = sum(w*y)/(sum(w)), time = min(time), sd = 1/sqrt(sum(w)), n = n()) %>% select(-labels)
  dfgroup$type = (dfgroup$n > 1)
  dfgroup$type[dfgroup$type] = "Molecule"
  dfgroup$type[dfgroup$type == F] = "Mix"
  
  if(keep.mix) {
    return(dfgroup)
  } else {
    return(filter(dfgroup, type != "Mix"))
  }
}
```
