## Supplementary material for "Correction of multiple-blinking artefacts in photoactivated localisation microscopy": Code and example data: Scoring.R

```
fix_clustering = function(labels) {
  wna = is.na(as.numeric(labels))
  labels = as.character(labels)
  if(length(wna) > 0) {labels[wna] = paste("N", 1:sum(wna))}
  
  wu = length(unique(labels))
  labels = factor(labels, labels = 1:wu)
  return(labels)
}

# Return the prior labeling probability
score_STC_prior = function(labels, pars, framerate) {
  m = moment.approx(pars[c(4,2,3)], 1/framerate)[1]
  N = length(labels)
  Nc = length(unique(labels))
  po = dnbinom(N-Nc, Nc, 1/m, log = T)
  return(po) 
}

score_STC_locs = function(locs, sds, xlims, ylims) {
  n = nrow(locs)
  
  vr = sum(1/sds^2)
  xtilde = c(sum(1/sds^2*locs[,1])/vr,sum(1/sds^2*locs[,2])/vr)
  
  #Location scoring
  ptscorex =
    (-(n-1)/2)*log(2*pi)-
    sum(log(sds))-
    1/2*sum(1/sds^2*(locs[,1]-xtilde[1])^2)-
    1/2*log(vr)+
    log(diff(pnorm(xlims, xtilde[1], sd = 1/sqrt(vr)) ))-
    log(diff(xlims))
  
  ptscorey =
    (-(n-1)/2)*log(2*pi)-
    sum(log(sds))-
    1/2*sum(1/sds^2*(locs[,2]-xtilde[2])^2)-
    1/2*log(vr)+
    log(diff(pnorm(ylims, xtilde[2], sd = 1/sqrt(vr)) ))-
    log(diff(ylims))
  
  ptscore = ptscorex+ptscorey
  
  return(ptscore)
}

score_STC_locs_update = function(locs, sds, mu.cen, mu.sig) {
  n = nrow(locs)
  M = length(mu.sig)
  
  vr = sum(1/sds^2)
  
  x = locs[,1]
  y = locs[,2]
  
  tilde = c(sum(1/sds^2*x)/vr,sum(1/sds^2*y)/vr)
  
  new.var = (mu.sig^2*vr+1)
  
  big.exp = 
    1/(new.var)*exp(-vr*(tilde[1]-mu.cen[,1])^2*1/(2*new.var))*exp(-vr*(tilde[2]-mu.cen[,2])^2*1/(2*new.var))
  
  #Location scoring
  ptscore = 
    -log(M)-
    n*log(2*pi)-
    2*sum(log(sds))-
    1/2*sum( ((x-tilde[1])/sds)^2  )-
    1/2*sum( ((y-tilde[2])/sds)^2  )+
    log(sum(big.exp))
  
  return(ptscore)
}

score_STC_time = function(time, pars, framerate, expEnd) {
  rF = pars[1]
  rD = pars[2]
  rR = pars[3]
  rB = pars[4]
  
  rO = rD+rB # Rate out of F
  pbleach = rB/rO # Probability of bleaching.
  
  time = as.integer(time)
  
  # Time scoring
  sigo = sort(unique(time))
  
  st = min(sigo)
  et = max(sigo)
  jumps = which(diff(sigo) > 1.5)
  
  nD = length(jumps)
  
  Fexits = sigo[c(jumps, length(sigo))]
  Fentry = sigo[c(1,jumps+1)]
  
  Flens = Fexits-Fentry
  Dlens = sigo[jumps+1]-sigo[jumps]
  
  uniform = (rF <= 0) # Fix for ramped starting times.
  if(!uniform) {
    ttscore = 
      dexp(st/framerate, rF, log = T)+
      sum(dexp(Flens/framerate, rD+rB, log = T))+
      sum(dexp(Dlens/framerate, rR, log = T))+nD*log(rD/(rD+rB))+
      log( rD/(rD+rB)*pexp((expEnd-et)/framerate, rR, lower.tail = F) + rB/(rB+rD) )
  } else {
    ttscore = 
      sum(dexp(Flens/framerate, rD+rB, log = T))+
      sum(dexp(Dlens/framerate, rR, log = T))+nD*log(rD/(rD+rB))+
      log( rD/(rD+rB)*pexp((expEnd-et)/framerate, rR, lower.tail = F) + rB/(rB+rD) )
  }
  
  
  return(ttscore)
}

# Score individual cluster.
score_STC_one = function(pts, sds, pars, noise, xlims, ylims, expStart, expEnd, framerate) {
  # Location scoring
  locs = pts[,1:2]
  ptscore = score_STC_locs(locs, sds, xlims, ylims)
  
  # Time scoring
  times = pts[,3]
  ttscore = score_STC_time(times, pars, framerate, expEnd)
  
  # total score
  noise_val = 1/( (expEnd-expStart)*diff(xlims)*diff(ylims))
  np = nrow(pts)
  nonoisescore = ptscore+ttscore
  
  if(np == 1) {
    out = log(exp(nonoisescore)*(1-noise) + noise*noise_val)
  } else {
    out = nonoisescore+log(1-noise)
  }
  
  return(out)
}

score_STC_one_update = function(pts, sds, pars, noise, expStart, expEnd, framerate, mu.cen, mu.sig) {
  # Location scoring
  locs = pts[,1:2]
  ptscore = score_STC_locs_update(locs, sds, mu.cen, mu.sig)
  
  # Time scoring
  times = pts[,3]
  ttscore = score_STC_time(times, pars, framerate, noise, expEnd)
  
  # total score
  return(ptscore+ttscore)
}

# Full scoring function
# Data should be provided with times in frames!
# pars  = c(lambda_OF, lambda_FD, lambda_DF, lambda_FB)
# noise = mean(noise intensity)/mean(total intensity)
# Framerate = framerate of camera.
score_STC = function(labels, pts, sds, pars, noise, framerate, expStart = 0) {
  ndf = data.frame(pts, sds)
  expEnd = max(pts[,3])
  
  spltdf = split.data.frame(ndf, labels)
  
  xl  = range(pts[,1]); yl = range(pts[,2])
  fl  = function(l) score_STC_one(l[,1:3], l[,4], pars, noise, xl, yl, expStart, expEnd, framerate)
  spc = sum(unlist(lapply(spltdf, fl)))
  
  return(spc)
}

score_STC_update = function(labels, pts, sds, pars, noise, framerate, expStart = 0, mu.cen, mu.sig) {
  ndf = data.frame(pts, sds)
  expEnd = max(pts[,3])
  
  spltdf = split.data.frame(ndf, labels)
  
  fl  = function(l) score_STC_one_update(l[,1:3], l[,4], pars, noise, expStart, expEnd, framerate, mu.cen, mu.sig)
  spc = sum(unlist(lapply(spltdf, fl)))
  
  return(spc)
}

# Generic function for scoring a given clustering.
score_clustering = function(labels, pts, score_function, ...) {
  labels_n = fix_clustering(labels)
  clus_score = score_function(labels = labels_n, pts = pts, ...)
  return(clus_score)
}

# Cluster data, and score.
cluster_and_score = function(pts, cluster_function, score_function, cluster.args = list(), score.args = list()) {
  # Clustering
  cluster.args[["pts"]] = pts
  clus  = do.call(cluster_function, cluster.args)
  
  # Scoring
  score.args[["pts"]] = pts
  score.args[["labels"]] = clus
  score.args[["score_function"]] = score_function
  score = do.call(score_clustering, score.args)
  
  # Results
  return(list(clustering = clus, score = score))
}
```
