## Supplementary material for "Correction of multiple-blinking artefacts in photoactivated localisation microscopy": Code and example data: STBHClus.R

```
require(fastcluster)
require(dplyr)
require(magrittr)
require(ClusterR)
require(dbscan)

# Naive clustering (Simple method)
# simple_clus = function(pts, Dspace, Dtime, dspace, dtime) {
#   pts$id = 1:nrow(pts)
#   pts = arrange(pts, time)
# 
#   untime = sort(unique(pts$time))
# 
#   Dspace[Dspace <= dspace] = 0
#   Dtime[Dtime <= dtime] = 0
# 
#   D = Dspace+Dtime
# 
#   ho = hclust(D, "single")
#   cc = cutree(ho, h = 0)
#   return(cc)
# }

simple_clus = function(pts, Dspace, Dtime, dspace, dtime) {
  pts$id = 1:nrow(pts)
  pts = arrange(pts, time)

  untime = sort(unique(pts$time))

  Dspace[Dspace > dspace] = 1e9
  Dtime[Dtime > dtime] = 1e9

  D = Dspace+Dtime

  ho = hclust(D, "single")
  cc = cutree(ho, h = 1e9-1)
  return(cc)
}

simple_clus_curve = function(pts, Dspace, Dtime, dspace, times) {
  Norig = nrow(pts)
  times = sort(setdiff(times, 0))
  nt = length(times)
  
  Nvals = numeric(nt+1); Nvals[1] = Norig
  tvals = c(0, times)
  
  for(i in 1:nt) {
    dtime = times[i]
    o = simple_clus(pts, Dspace, Dtime, dspace, dtime)
    Nvals[i+1] = length(unique(o))
  }
  
  return(cbind(tvals, Nvals))
}

simple_curve_to_pars = function(curveout) {
  pf = function(par) {
    par = abs(par)
    n = par[1]; to = par[3]; nb = par[2]
    oo = n*(1+nb*exp((1-curveout[-1,1])/to))
    rt = mean( (curveout[-1,2]-oo)^2 )
    return(rt)
  }
  
  hi = boptim.try(c(3,10), pf, n = 100, plt = F)
  return(abs(hi$par))
}

simple_clus_extractN = function(pts, Dspace, Dtime, dspace, N, t0) {
  g1 = simple_clus(pts, Dspace, Dtime, dspace, t0)
                   
  N0 = length(unique(g1))
  if(N0 < N) {
    sgn = -1
  } else {
    sgn = 1
  }
  
  sgc = sgn
  i = 1
  while(sgc == sgn) {
    g1 = simple_clus(pts, Dspace, Dtime, dspace, t0+2*sgn*i)
    i = i+1
    N0 = length(unique(g1))
    overprint(N0)
    sgn = -sign(N-N0)
  }
  
  return(g1)
}

# Cluster into a fixed number of partitions. Use this.
STBHClus_N = function(pts, N, D, method = "ward.D2") {
  prop = fastcluster::hclust(D, method = method)
  prop = cutree(prop, k = N)
  return(prop)
}

# Code below not used.
STBHClus_base = function(df, ds, dt) {
  locs = df[,1:2]
  time = as.matrix(df[,3])
  
  tr = diff(range(time))
  D1 = dist(locs)
  D2 = dist(time)
  D2[D2 > dt] = tr
  D2[D2 <= dt] = 0
  
  D = D1+D2
  prop = fastcluster::hclust(D, method = "complete")
  prop = cutree(prop, h = ds)
  return(prop)
}

minpo = function(v) {
  v[v < 0] = Inf
  return(min(v))
}

# 'Stitch' together a clustering based on a subdivised region.
stitch_clustering = function(df, ds, dt, clus, grid, stitch.d, plt = F) {
  locs = df[,1:2]
  df$row = 1:nrow(df)
  
  gridr = grid[,2:(ncol(grid)-1)]
  if(is.null(dim(gridr))) {gridr = t(t(gridr))}
  
  df$clus = clus
  
  locs.c = data.frame(locs, clus)
  o = locs.c %>% group_by(clus) %>% summarise(
    xl = minpo(gridr[1,]-max(x)), xr = minpo(min(x)-gridr[1,]),
    yt = minpo(min(y)-gridr[2,]), yb = minpo(gridr[2,]-max(y)))
  o[o < 0] = 2*stitch.d
  
  o = o %>% group_by(clus) %>% summarise(xd = min(xl,xr), yd = min(yt, yb), keep = min(xd,yd) <= stitch.d) %>% select(clus, keep) %>% filter(keep == T)
  
  df.keep   = df %>% filter(not(clus %in% o$clus))
  df.stitch = df %>% filter(clus %in% o$clus)
  
  stclus = STBHClus_base(df.stitch[,1:3], ds, dt)
  df.stitch$clus = paste(stclus, "st", sep = "")
  
  odf = rbind(df.keep, df.stitch)
  odf$clus = as.numeric(factor(odf$clus))
  
  if(plt) {
    plot(df[,1:2], asp = 1, type = "n")
    abline(h = gridr[2,], v = gridr[1,])
    abline(h = c(gridr[2,]-stitch.d, gridr[2,]+stitch.d), v = c(gridr[1,]-stitch.d,gridr[1,]+stitch.d), lty = 2, col = "red")
    df.keep %$% points(.[,1:2], pch = 19, col = "lightgray")
    df.stitch %$% points(.[,1:2], pch = 19, col = factor(df.stitch$clus))
  }
  
  odf = arrange(odf, odf$row)
  
  return(odf)
}

#### Spatio-temporal clustering algorithm for large datasets
# df       : dataframe with columns 'x', 'y', 'time'. Time should be in seconds.
# ds       : distance in space within which to look for cluster members.
# dt       : distance in time within which to look for cluster members.
# N        : number of cuts per axis for subdivision of data.
# stitch.d : distance from grid within which to stich together subregion clustering results.
STBHClus = function(pts, ds, dt, N = 4, stitch = T, stitch.d = ds/2, plt = F) {
  df = pts
  df$row = 1:nrow(df)
  boundaries = rbind(range(df[,1]), range(df[,2]))
  
  cx = seq(boundaries[1,1], boundaries[1,2], length.out = N+1)
  cy = seq(boundaries[2,1], boundaries[2,2], length.out = N+1)
  
  if(plt) {
    plot(df[,1:2], asp = 1, pch = 19, col = "gray", type = "n")
    abline(h = cy, v = cx, lty = 2, col = "red")
    abline(h = c(cy-stitch.d, cy+stitch.d), v = c(cx-stitch.d,cx+stitch.d), lty = 3, col = "yellow")
  }
  
  M = 1
  clustering = c()
  dfo = data.frame()
  for(i in 1:N) {
    for(j in 1:N) {
      st = paste(i,"x",j, sep = "")
      dfs = subset(df, dplyr::between(df$x, cx[i], cx[i+1]) & dplyr::between(df$y, cy[j], cy[j+1]))
      cl = STBHClus_base(dfs[,1:3], ds, dt)
      cl = paste(cl, "-", M, sep = "")
      dfs$clus = cl
      
      dfo = rbind(dfo, dfs)
      
      M = M+1
      if(plt) {points(dfs$y~dfs$x, pch = 19, col = factor(cl) )}
    }
  }
  
  dfo$clus = factor(dfo$clus)
  dfo$clus = as.numeric(dfo$clus)
  dfo = arrange(dfo, dfo$row)
  
  if(stitch) {
    odf = stitch_clustering(dfo[,1:3], ds, dt, dfo[,5], rbind(cx,cy), stitch.d = stitch.d, plt = plt)
  } else {
    odf = dfo
  }
  
  return(odf$clus)
}

# Simple Kmeans-based clustering.
ST.Kmeans = function(pts, k, s, nrestart = 20, maxiter = 10) {
  pts2 = pts
  pts2$time = pts2$time*s
  ko = kmeans(pts2, k, iter.max = maxiter, nstart = nrestart)
  return(ko$cluster)
}
```
