## Supplementary material for "Correction of multiple-blinking artefacts in photoactivated localisation microscopy": Code and example data: main.R

```
source(file.path("Dependencies", "Dependencies.R"))

MBC = function(data, parameters = parameters) {
  # Data is a data-frame with columns (X, Y, time, localisation uncertainty standard deviation)
  framerate = parameters$framerate
  alpha = parameters$alpha
  nopt = parameters$nopt
  nu = parameters$nu
  cut = parameters$cut
  t.thres = parameters$t.thres
  
  X = ppp(x = data[,1], y = data[,2], marks = as.integer(data[,3]), window = owin(range(data[,1]), range(data[,2])))
  unc = data[,4]
  
  rate.estimates = estimate.model(X, framerate, unc, alpha = alpha , cut = cut, nopt = nopt, nu = nu)
  ord.params = as.numeric(rate.estimates$par[c(4,2,3,1)])
  
  eint = (1-alpha)*intensity(X)
  declus = STBHClus_optimize(data, framerate, ord.params, eint, t.thres)
  
  MBC.data = declus$merged %>% select(-time)
  MBC.data = rename(MBC.data, uncertainty = sd)
  
  return(MBC.data)
}
```
