## Supplementary material for "Correction of multiple-blinking artefacts in photoactivated localisation microscopy": Code and example data: README.txt

# Multiple-blinking correction

This is the R code for the paper: **Correction of multiple-blinking artefacts in photoactivated localisation microscopy**

The code takes as input data in a Comma Separated Value file (.csv) with the fields (x,y,time,sd) representing x, y, frame number and localisation uncertainty (standard deviation) respectively.

We provide two examples, a set of fluorophore proteins on a grid, with either light or heavy blinking (Data_light.csv, Data_heavy.csv). The code outputs a set of corrected localisations, obtained after estimating blinking dynamics, and clustering input localisations accordingly.

## Coding environment
We assume that RStudio and R are installed (tested on R version 4.0.4), as well as the following R packages:
- tidyverse
- magrittr
- wrapr
- spatstat
- multiway
- fastcluster
- dbscan

1. Extract the `mbc.zip` archive 
2. Place data to be processed in this folder (e.g. Data_light.csv)
3. In RStudio, open Run.R
4. Set the working directory (line 1) to the extracted folder (the folder that contains main.R). Remember that for Windows operating systems file paths us / not \
5. Set the data file to be processed (line 3) e.g. data = read.csv("Data_light.csv", )
6. Set the camera frame rate (line 6) measured in frames per second (default = 25)
7. Set the expected fraction of non-noise points (alpha, line 7, 1 by default)
8. Run `source("Run.R")` 

Code will perform the multiple-blinking correction. A .csv file containing the fields (x,y,sd,n) representing corrected x, y coordinates, updated localisation uncertainty (standard deviation) and number of input points merged to one, corrected localisation will be saved.
