## Supplementary material for "Correction of multiple-blinking artefacts in photoactivated localisation microscopy": Code and example data: Run.R

```
setwd("enter-folder-path")
source("main.R")
data = read.csv("enter-filename-to-process", )

parameters = list(
  framerate = 25,
  alpha = 1,
  

  
  cut = NA,
  nopt = 15,
  nu = 50,
  t.thres = seq(0.03, 0.25, length.out = 50)/100
)


corrected = MBC(data, parameters = parameters)

write.csv(corrected, "Data_corrected.csv", row.names = F, quote = F)

# Plotting data before and after MBC correction.
par(mfrow = c(1,2), mai = rep(0.4,4))
plot(data[,1:2], pch = 19, cex = 1, asp = 1, main = "Input data")
plot(corrected[,1:2], pch = 19, cex = 1, asp = 1, main = "Corrected data")
```
